# Global Scientific and Public Interest in Birds and their Conservation

**DOI:** 10.1101/2024.12.12.628088

**Authors:** Haozhong Si, Stuart H. M. Butchart, Fan Yu, Changjian Fu, Zhongqiu Li, Enrico Di Minin

## Abstract

Greater public support to conservation is needed to help address the global biodiversity crisis. However, human attention towards biodiversity is taxonomically biased, leaving many species neglected and their conservation needs unmet. Here we investigated the drivers of public and scientific interest for 10,509 bird species, measured by Wikipedia pageviews and articles in Web of Knowledge. We then used a quasi-experimental design to test the effect on public interest of published updates to the International Union for the Conservation of Nature (IUCN) Red List of threatened species (N = 479). Public and scientific interest were strongly predicted by species’ charisma, familiarity, utilization, and Red List category. While threatened species received significantly more attention, IUCN Red List category revisions produced asymmetric public response: uplistings (to categories of higher extinction risk) generated a significant but modest immediate increase in pageviews, while downlistings (to categories of lower risk) showed no detectable effect. The traits that predicted responsiveness, genuineness of change (real biological shifts rather than improved knowledge or revised taxonomy) and increasing population trend, differed from those driving underlying interest. These findings underscore the need to raise the profile of species lacking the charismatic or familiar traits that typically attract public and scientific interest.

## Introduction

The ongoing global biodiversity crisis is also affecting ecosystem functioning and human well-being ^1^. The threats to global biodiversity, for example land use change, are well-documented, yet slowing the current rates of biodiversity loss remains a challenge ^2–4^. Conservation efforts often target a limited set of taxa, including charismatic species or those that are of local concern rather than globally threatened, while the majority of species in need of conservation attention still receive insufficient action^5^. To advance more balanced conservation efforts, it is necessary to increase public awareness and support for a broader set of species ^6^. The Kunming-Montreal Global Biodiversity Framework (GBF) aims to engage all of society to halt and reverse biodiversity loss. Improving public awareness and accounting for a diverse set of values of species can encourage more just and favorable attitudes towards biodiversity conservation ^7^.

Although less studied and seldom measured, investigating the societal and scientific interests in biodiversity can help better direct limited resources to species that are most in need of conservation actions ^8–11^. It is well known that species that attract strong public interest are more likely to receive funds and mobilize support for their conservation ^5^. Charismatic animals, such as tigers (*Panthera tigris*), gain significant attention from both the public and researchers ^12,13^, whereas species in other groups, such as insects, receive far less attention and conservation support, despite their essential role in ecosystems ^14,15^. This bias poses serious risks for species that need conservation but fail to attract sufficient attention and conservation resources ^11,16^. Societal preferences can also influence scientific research, leading scientists to prioritize studying larger, more charismatic species regardless of their conservation status ^17^. Important and enduring taxonomic biases exist in the species that have been studied, influencing both the quantity and quality of scientific knowledge available for different species ^18^. The long-term success of conservation efforts hinges on the availability of a robust body of evidence-based research about all species or habitats in need of conservation ^19^.

Disparities in societal and scientific knowledge of species can be influenced by various factors. Phenotypic traits, such as body size and coloration, directly influence species’ charisma and feasibility for research ^20–22^. Large mammals ^12,20^ or colorful birds ^23–25^, butterflies ^26^ and plants ^22^ are more likely to be appreciated and studied, while smaller or cryptic species, like many insects ^14,15^, are frequently overlooked and typically studied less. Species with larger populations and distributions are also more likely to be studied and recognized by the public ^27,28^. Conversely, geographically restricted species, often found in regions with low human development indices, receive less attention than widely distributed species ^29^. Moreover, cultural factors can increase public attention and conservation efforts, such as for species used as mascots like the bald eagle (*Haliaeetus leucocephalus*) ^30^. The role of a species’ conservation status in influencing levels of attention is particularly complicated. While it has been suggested that a species being listed as threatened can attract attention through conservation efforts and media publicity ^27,31,32^, many threatened plant and insect species remain overlooked due to a lack of familiarity or appeal ^10,13,15,22,33^. A comprehensive study investigating what factors shape both public and scientific interest in bird species worldwide is still missing. Existing studies have focused on a narrow set of static factors ^27^ or limited regions ^34–36^, lacking a comprehensive global analysis that considers a broader range of influences.

In this study, we investigated the global drivers of public and scientific interest in bird species (Figure 1). Birds are an ideal group for this analysis due to their cultural visibility and the comprehensive data available on their traits and conservation status ^37–41^. First, we used Generalized Linear Mixed Models (GLMMs) to understand what factors affect scientific and public interest, measured respectively by the number of articles mentioning each species indexed in the Web of Knowledge (WOK) and the number of relevant Wikipedia page views across all available languages respectively. We then used a Gaussian Linear Mixed Model to analyze the relative bias between these two metrics of attention. Finally, we employed a quasi-experimental Difference-in-Differences (DiD) design to test the causal impact of IUCN Red List category changes on public interest. See Supplementary Methods for more details.

**Figure 1.**
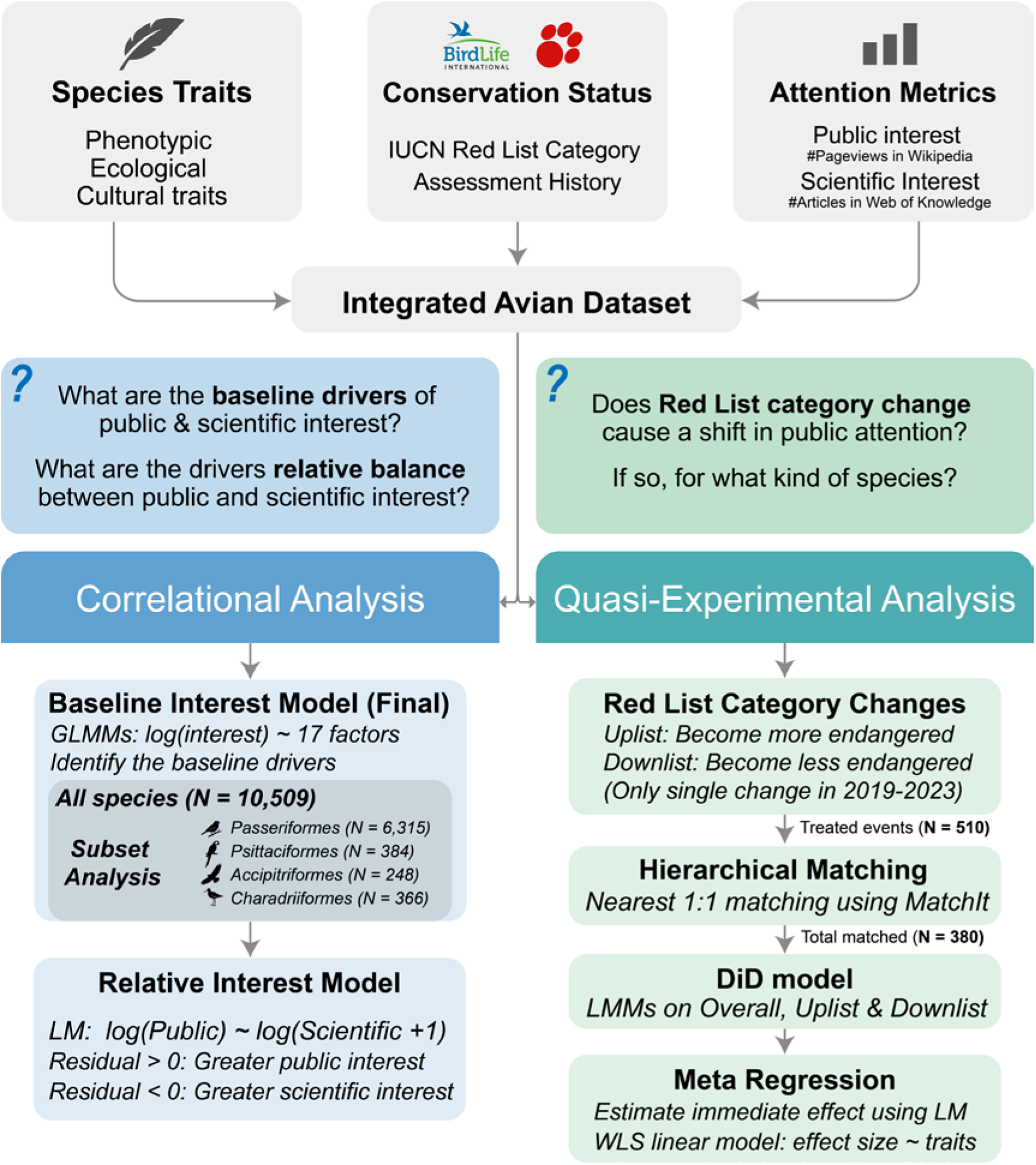
Conceptual flowchart of the analytical methodology. The flowchart illustrates the study’s two main analytical streams. The Correlational Analysis (left, blue) uses static, species-level data and Generalized Linear Mixed Models (GLMMs) to identify the baseline drivers of public and scientific interest, as well as the drivers of the relative balance between them. The Quasi-Experimental Analysis (right, green) uses time-series data and a Difference-in-Differences (DiD) design to test the causal impact of IUCN Red List category changes. This analysis involves a multi-stage hierarchical matching procedure to create a robust control group (N = 380), followed by a series of Linear Mixed-Effects Models (LMMs) to estimate immediate and sustained treatment effects for the overall, uplist, and downlist samples. A final meta-regression was then used to investigate what species traits predict a stronger public response.

## Results

### Distribution of scientific and public interest

Overall, the scientific names of 696 bird species (i.e., 6.22% of the 11185 species, 2.4% were threatened) were not mentioned in any scientific article in Web of Knowledge. For Web of Science, 3841 species were not mentioned (i.e., 31.12% of the 11185 species, 7.3% were threatened). In contrast, only 238 bird species (i.e., 2.13% of the 11185 species, 8.0% were threatened) lacked their own Wikipedia pages, mostly due to differences between the taxonomic systems used by Wikipedia contributors and the IUCN/BirdLife checklist. A full list of species missing from either source is provided in Supplementary Data 1.

The number of scientific papers mentioning each bird species varied by four orders of magnitude and exhibited a skewed distribution (median ± SE = 8 ± 15.78, range = 0 – 174455) (Figure 2). Wikipedia pageviews, instead, were more evenly distributed (median ± SE = 69831 ± 11281.16; range = 714– 24530989) (Figure 2). Despite different distributions, the two metrics were strongly correlated (Adjusted R² = 0.626, F_(1, 10507)_ = 17590, p < 0.001).

**Figure 2.**
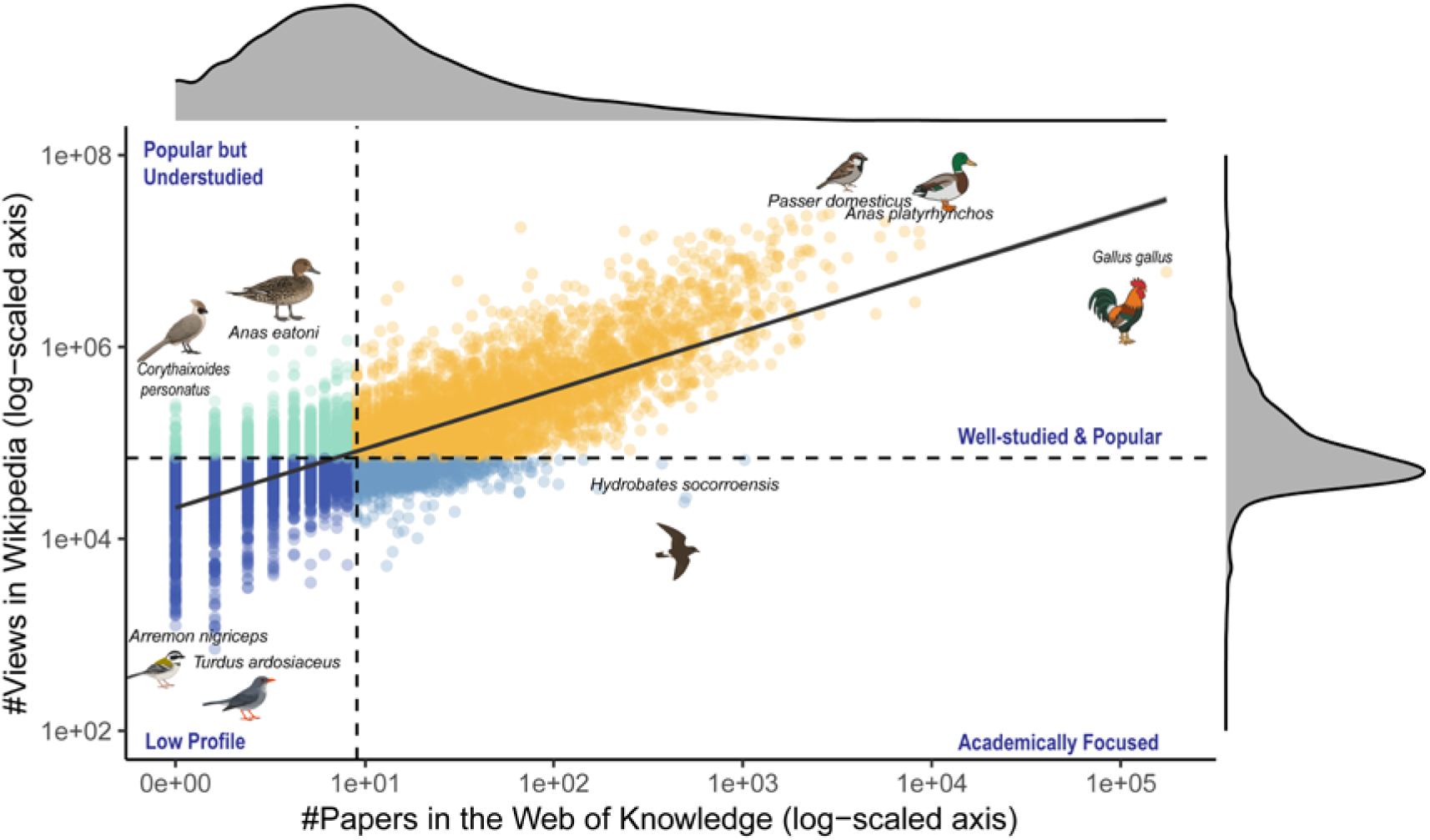
Relationship between public and scientific interest across bird species. Each point represents a single species (N = 10509), plotted by its scientific interest (total number of Web of Knowledge papers, x-axis) and public interest (total Wikipedia pageviews, y-axis). Both axes are log-scaled to facilitate visualization. The solid black line shows the overall positive correlation, fitted using a simple linear model on the log-transformed data (F_(1, 10507)_ = 17590, Adj. R^2^ = 0.626, p < 0.001). The residual (vertical distance of a point from this line) indicates its attention bias. Positive residual indicates greater public interest than predicted by their scientific interest, negative residual indicates greater scientific interest. To better illustrate these patterns, the plot is divided into four conceptual regions based on median values of interest and the regression line. Species with minimal public and scientific interest are marked as ‘Low Profile’. Those with high public but low scientific interest are ‘Popular but Understudied’. For well-studied species, those with high public interest are ‘Well-studied & Popular’, while those with low relative public interest are ‘Academically Focused’. Representative species are labeled to illustrate each quadrant. Marginal density plots show the highly skewed distribution of both scientific (top) and public (right) interest.

### Drivers of baseline scientific and public interest

Our GLMMs successfully identified a consistent set of traits shaping both public and scientific interest. The models demonstrated strong explanatory power, with the selected fixed effects accounting for a substantial proportion of the variance in both public (Marginal R² = 0.585) and scientific (Marginal R² = 0.609) interest. However, the relative importance of these drivers differed between the two domains (Figure 3).

**Figure 3.**
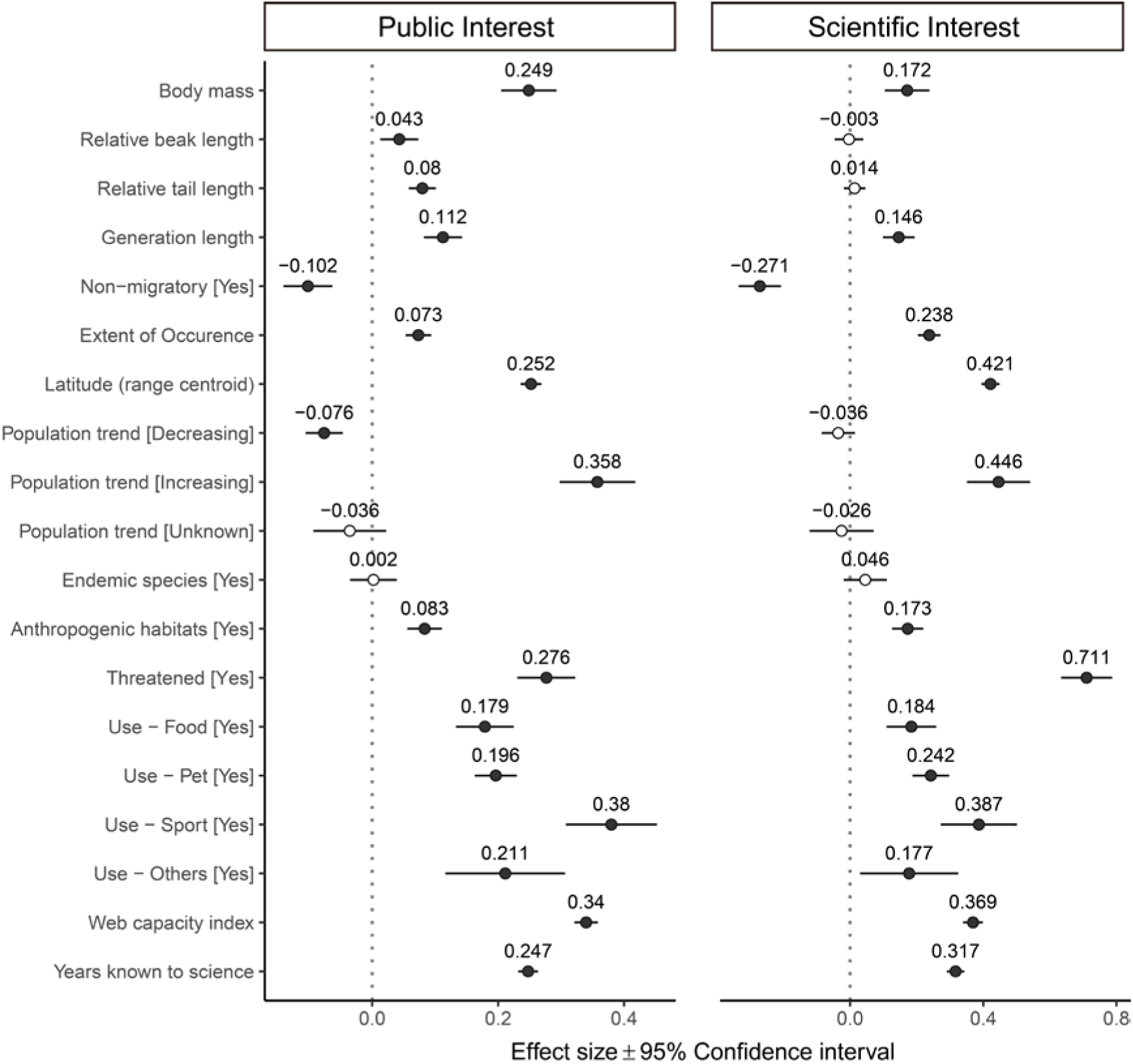
Drivers of scientific and public interest across bird species. Forest plots summarize the estimated parameters (effect size mean ± 95% CI, N = 10509) based on (truncated for scientific interest model) negative binomial generalized linear mixed models (Eq. 1). Baseline levels for multilevel factor variables are: Population trend (Stable). The left one modelled public interest (marginal R^2^ = 0.585), and the right one modelled scientific interest (marginal R^2^ = 0.609). Filled points indicate statistically significant effects (p < 0.05). Estimated regression parameters and p-values are in Table S1.

Larger body mass, a trait typically linked to species charisma, was strongly associated with greater interest in both the public (Estimate ± SE: 0.25 ± 0.02, p < 0.001) and scientific (0.17 ± 0.03, p < 0.001) interest models. Factors related to familiarity consistently emerged as strong predictors of public interest. Species that have been known to science for longer (Public: 0.28 ± 0.01; Scientific: 0.31 ± 0.01, both p < 0.001) and those occurring in regions with a higher web capacity index (the proportion of global internet users in countries across a species’ native range; Public: 0.34 ± 0.01; Scientific: 0.37 ± 0.02, both p < 0.001) received significantly greater interest. Similarly, birds occurring in anthropogenic habitats (Public: 0.08 ± 0.01; Scientific: 0.17 ± 0.02, both p < 0.001) attracted significantly more interest. The consumptive use of species was also a major predictor of interest. Birds used and traded for food (Public: 0.18 ± 0.04; Scientific: 0.18 ± 0.02, both p < 0.001), pets (Public: 0.20 ± 0.02; Scientific: 0.24 ± 0.03, both p < 0.001), sport hunting (Public: 0.38 ± 0.04; Scientific: 0.39 ± 0.06, both p < 0.001) and other use categories (Public: 0.18 ± 0.08, p < 0.001; Scientific: 0.18 ± 0.08, p = 0.018) were among the most popular across both domains.

Conservation-related attributes also showed a significant influence. Both public and scientific interest were significantly higher for species with ranges centered at higher latitudes (Public: 0.25 ± 0.01; Scientific: 0.42 ± 0.01, both p < 0.001) and those with larger extent of occurrence (Public: 0.07 ± 0.01; Scientific: 0.23 ± 0.02, both p < 0.001). A species’ IUCN Red List category was a strong predictor, with threatened species receiving significantly more interest than non-threatened (LC and NT) species for both public (0.28 ± 0.02, p < 0.001) and scientific (0.71 ± 0.04, p < 0.001) interest. Similarly, species with increasing population trends were highly prominent in both domains (Public: 0.36 ± 0.03; Scientific: 0.45 ± 0.05, both p < 0.001). Interestingly, a decreasing population trend was associated with lower public interest (−0.08 ± 0.02, p < 0.001) but had no significant effect on scientific interest (p = 0.16).

To test the generality of our findings, we repeated the baseline interest analyses on four major avian orders with distinct ecological and cultural profiles: Passeriformes (N = 6315, the largest avian order), Psittaciformes (parrots, N = 384), Accipitriformes (most raptors, N = 248), and Charadriiformes (shorebirds, gulls and auks, N = 366). Several key factors of attention were consistent across these diverse clades (Figure S1-2, Table S2). Factors related to familiarity, such as more years known to science, a higher web capacity index and ranges at higher latitudes, emerged as universally strong and positive predictors for both public and scientific interest in nearly all analyzed orders. Similarly, an increasing population trend was also a consistent correlate of higher attention. Beyond these shared drivers, some patterns were specific to each domain (see Appendix Results 1 and Figures S3-6).

### Bias between scientific and public Interest

Our analysis of the relative balance between public and scientific interest revealed a clear and significant profile of traits that disproportionately attracted attention from the public (Figure S7, Table S3). The model successfully explained a modest, but statistically significant portion of the variance (Marginal R² = 0.094). Species with a larger body mass (Estimate ± SE: 0.13 ± 0.02, p < 0.01), longer beak (0.03 ± 0.01, p = 0.02) and tail (0.05 ± 0.01, p < 0.01) were significantly more popular with the public than with scientists. Traits related to human familiarity also attracted significantly more public than scientific interest. Species utilized for food (0.04 ± 0.02, p = 0.04), pet (0.05 ± 0.02, p < 0.01) or sport hunting (0.09 ± 0.03, p < 0.01) were all significantly more popular with the public than expected from their scientific interest. A higher web capacity index (0.08 ± 0.01, p < 0.01) and more years known to science (0.08 ± 0.01, p < 0.01) also predicted relatively more public interest. Species with an increasing population trend also tended to attract relatively more public interest (−0.07 ± 0.03, p = 0.01). In contrast, threatened species (−0.14 ± 0.02, p < 0.01) and those with a decreasing population trend (−0.05 ± 0.01, p < 0.01) tended to attract relatively more scientific interest.

### Response to IUCN Red List category changes

Species are regularly reassessed for the IUCN Red List, which in some cases leads to their reclassification into a higher or lower Red List category, either because of genuine improvement/deterioration in status, or as a result of improved knowledge or taxonomic revisions. To isolate the causal effect of IUCN Red List category changes from background trends, we compared Wikipedia pageview trajectories of treated species to matched controls using a Difference-in-Differences design. A significant positive background trend in pageviews was detected for all species in the post-event period (Estimate ± SE: 0.08 ± 0.02, p < 0.01) (Figure 4). After accounting for this trend, uplistings showed a significant immediate increase in pageviews relative to matched controls (0.067 ± 0.003, p = 0.02) (Figure 5a). No significant immediate effect was found for downlistings (p = 0.49), nor did either group show a significant change in long-term trend (uplist: p = 0.86; downlist: p = 0.24).

**Figure 4.**
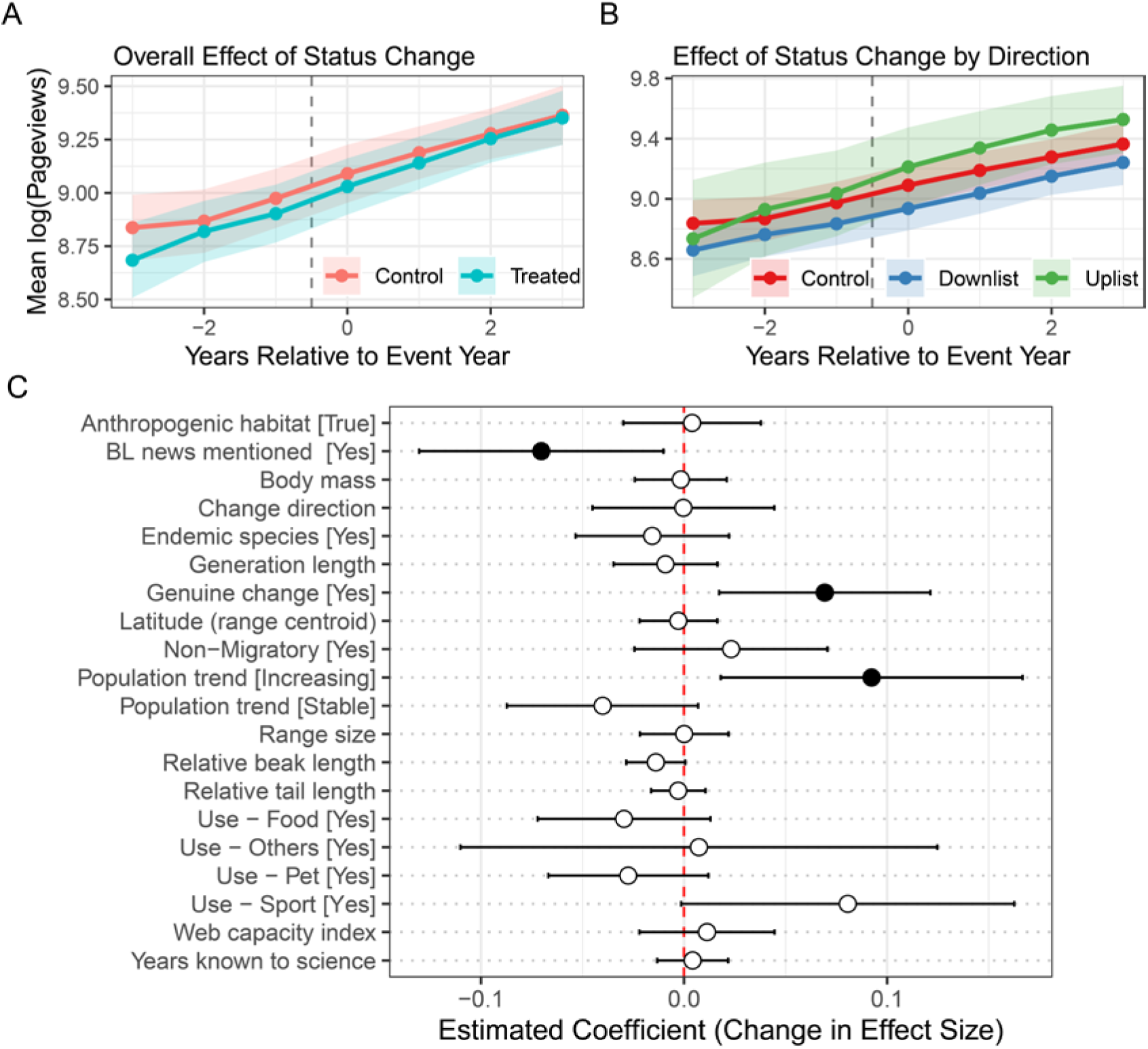
Public responsiveness to IUCN Red List category changes and the predictable sensitivity to uplist events. (A) The overall mean trend in log-transformed Wikipedia pageviews for treated species (those with a Red List category change, blue) and their matched controls (red). The parallel trends before the event (Year 0, indicated by the dashed line) and the lack of divergence after the event visually demonstrate the null average effect of a status change. Shaded areas represent 95% confidence intervals. (B) Mean pageview trends for the control group (red) versus the uplist (green) and downlist (blue) subsets. (C) Coefficient plot of the meta-regression model identifying species traits that predict public response to Red List Category changes (F_(20, 359)_ = 2.13, p = 0.004; Adj. R² = 0.056). “BL news mentioned” refers to category changes mentioned in BirdLife International’s annual Red List news update, included as a control variable. Each point represents the estimated coefficient from the model, with horizontal lines showing the 95% confidence interval. Filled points indicate statistically significant effects (p < 0.05). Estimated regression parameters and p-values are in Table S5.

**Figure 5.**
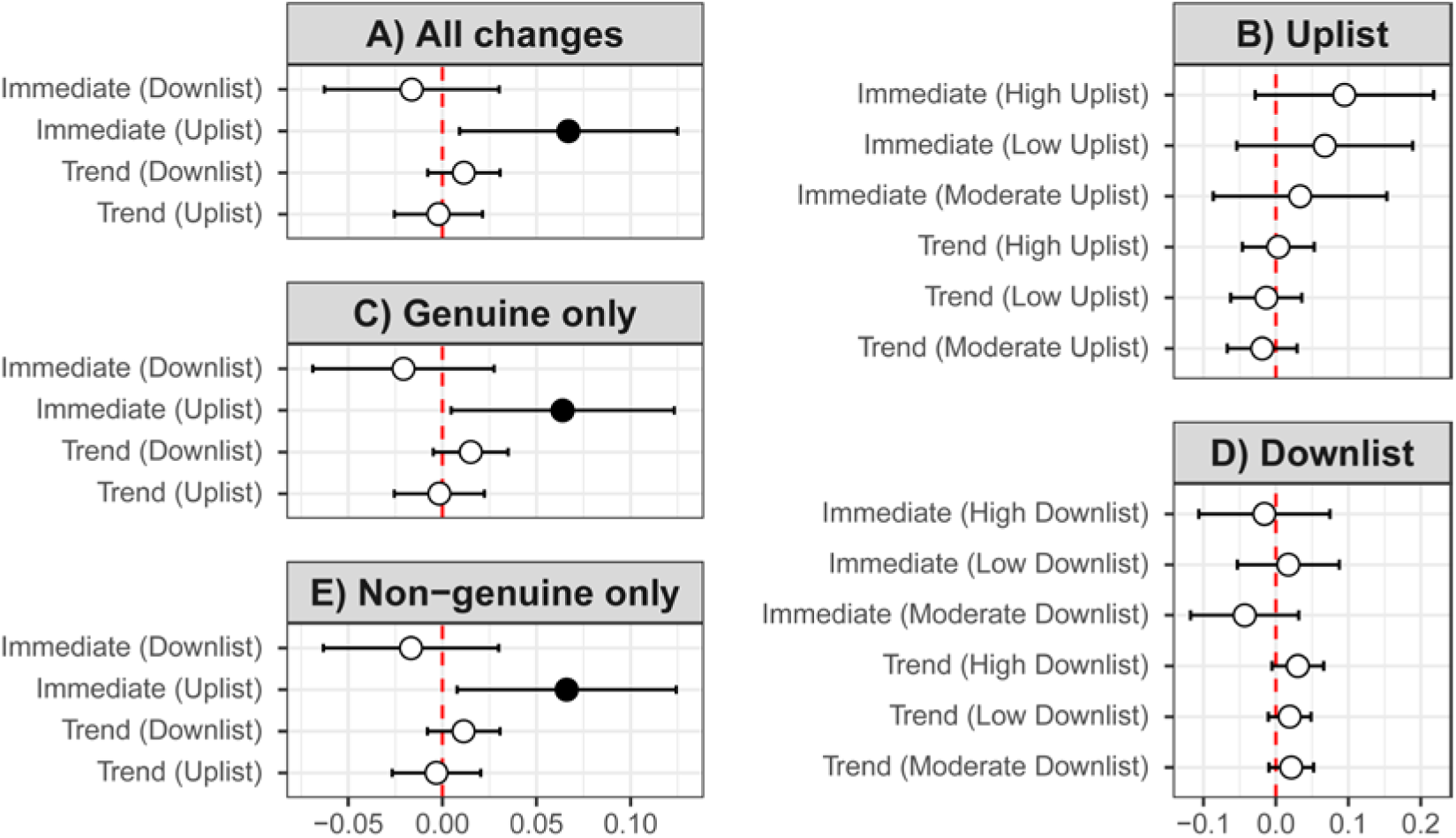
Difference-in-Differences estimates of the effect of IUCN Red List category changes on public interest. Fxed-effect coefficients and 95% confidence intervals from three separate Linear Mixed-Effects Models. (A) The primary model, showing the average immediate and sustained effects for all uplist and downlist events relative to a matched control group. (B) The sub-analysis model for uplist events, showing the effects for low, moderate, and high magnitude changes relative to their matched controls. (C) The primary model restricted to treated species with genuine status changes only (n = 108), using the same matched pairs as panel A. (D) The sub-analysis model for downlist events, showing the effects for low, moderate, and high magnitude changes relative to their matched controls. (E) The primary model restricted to treated species with non-genuine status changes only (n = 272), using the same matched pairs as panel A. Panels A, C, and E use the same covariate specification (the eight matching variables) and differ only in the treated pool. “Immediate Effect” corresponds to the “post event” interaction term, while “Trend Effect” corresponds to the “time since event” interaction term. Filled points indicate statistically significant effects (p < 0.05). Full model results are provided in Table S4.

Focusing on the uplisting and downlisting subsets, we further analyzed the public response in relation to the level of extinction risk. Neither an immediate nor a sustained effect was detected for uplistings to categories of low risk (NT), moderate risk (VU), or high risk (EN or CR), nor downlistings from categories of low risk (NT), moderate risk (VU), or high risk (EN or CR) (Figure 5b, 5d).

To further test whether the public response depended on the genuineness of category change, we split the matched pairs into genuine (real changes to the extinction risk) or non-genuine (revision due to improved knowledge or revised taxonomy) subsets and re-ran the DiD models on each (Figure 5c, 6e). The immediate uplist effect was significant and of nearly identical magnitude in both the genuine-only subset (0.06, p = 0.03) and the non-genuine-only subset (0.07, p = 0.03), each closely matching the pooled estimate (0.07, p = 0.02). This means even non-genuine uplistings trigger a public response comparable with genuine ones. In contrast, downlistings showed no significant effect in either subset.

To understand what affects public response to IUCN Red List category changes, we conducted a meta-regression analysis to identify the traits predicting the magnitude of the immediate attention shift. The full meta-regression model, pooling all treated species, was statistically significant (Overall model F_(20, 359)_ = 2.13, p = 0.004; Adj. R² = 0.056). The genuineness of change emerged as a significant predictor (Figure 4c), where species with genuine Red List category change showed a significantly larger immediate pageview response than those with non-genuine changes (Estimate ± SE: 0.07 ± 0.03, p < 0.01).

## Discussion

Our study revealed a set of global drivers of public and scientific interest in bird species, with substantial overlap but also notable divergences between the two. Interest is largely shaped by species charisma, familiarity, direct utility or relationships with humans, and conservation-related attributes. While higher risk of extinction, based on the IUCN Red List category, strongly predicted greater public interest, our quasi-experimental analysis found that public responses to category changes were modest and asymmetric. Uplistings produced a significant immediate increase in Wikipedia pageviews relative to matched controls, whereas downlistings showed no detectable immediate effect. Changes in Red List category did not lead to a significant shift in long-term pageview trends, nor significant effects at different magnitude levels. Genuine and non-genuine uplistings generated comparable immediate increases, yet genuineness of change was the strongest predictor of response magnitude in a meta-regression across species.

Regarding the drivers of baseline interest, morphological traits associated with charisma, especially body mass, were strong and consistent predictors of public interest. Larger body mass was associated with greater interest in both domains, aligning with evidence that larger species are more noticeable and perceived as charismatic ^20,25,31^. Relative beak and tail length also attracted significantly more public interest, yet neither predicted scientific attention. This public bias was confirmed by the relative interest model, where all three traits significantly favored public over scientific attention. Across the four subset orders (see Supplementary Results and Figures S3-S6), body mass predicted public interest in all groups, but its effect on scientific interest was limited to Charadriiformes and, marginally, Accipitriformes. It was non-significant for Passeriformes and Psittaciformes. Among Psittaciformes, relative tail length rather than body mass was the dominant morphological predictor of public interest, consistent with the visual prominence of elongated tails in this group ^42^. These clade-specific patterns suggest that the traits driving perceived charisma vary across bird groups, and that scientific interest is generally less responsive to morphological appeal than public interest. Charismatic species with striking traits or cultural associations often attract more attention and conservation funding ^13^. While beneficial for these species, this bias disadvantages less charismatic but equally threatened taxa ^13,36^, underscoring the need to mitigate charisma-driven imbalances in conservation attention.

Traits related to familiarity—through cultural association and ecological prevalence—is a major driver of both public and scientific interest. Familiarity correlates with internet popularity ^27,30,43^ and scientific interest ^22^. Species used as food, kept as pets, or hunted for sport consistently receive greater interest, possibly reflecting relevant cultural traditions such as aviculture and hunting ^44,45^.

Species with large range sizes and those at higher absolute latitudes attracted more public and scientific interest. Given that Wikipedia usage and research institutions are concentrated in Europe and North America ^46,47^, the latitude effect likely reflects geographic biases in internet access and research capacity. The positive association with the Web Capacity Index supports this interpretation ^27,48^. Historical factors also play a role: species named earlier tend to receive more attention, reflecting temporal biases in scientific recognition and cultural prominence ^49,50^. Conversely, species with smaller ranges and lower latitudes receive less attention. As bird diversity peaks near the equator ^51^, this concentration of interest on temperate species familiar to internet users and researchers leaves many tropical species under-studied and receiving inadequate conservation responses. Addressing the bias in scientific interest is essential for effective and comprehensive biodiversity conservation and long-term sustainability ^16,52^.

Beyond familiarity and utility, current Red List category was a strong predictor of interest. Threatened species receive significantly more attention globally, even after controlling for confounding traits, consistent with findings in other taxa ^27,31,32^. This pattern held across three of our subset bird orders: Passeriformes, Accipitriformes, and Psittaciformes. Charadriiformes was the exception, where Red List category did not significantly predict public interest and its effect on scientific interest was less pronounced, suggesting that the influence of Red List category on public attention varies across clades.

It could reasonably be expected that if a species is uplisted to a higher extinction risk category, this should cause a corresponding increase in public attention ^53^. Our DiD analysis, however, found only a modest immediate increase for uplistings, with no significant sustained effects and no detectable response to downlistings or to changes at individual magnitude levels. That pattern is also perhaps unsurprising given that large numbers of category revisions (∼100 per year) are published by BirdLife International (and reflected on the IUCN Red List) each year, and that only a handful are proactively highlighted in the associated news releases and media coverage. Nonetheless, while the IUCN Red List is critical for conservation prioritization ^54,55^, Red List category changes do not automatically translate into public awareness.

The factors that predicted the strength of public response to Red List category changes were selective and differed notably from the key predictors of baseline interest. The meta-regression identified genuineness of change as the strongest predictor of response magnitude, where genuine changes generate larger pageview increases than non-genuine ones. Increasing population trend also predicted a stronger response. In contrast, traits related to familiarity, charisma, and direct utilization—all strong predictors of baseline interest, showed no significant effect on response magnitude. This suggests that the traits that attract public attention in general are not the same as those that amplify attention when species’ Red List categories are revised. Notably, while genuineness of change predicted larger effect sizes across species in the meta-regression, the DiD analysis found that genuine and non-genuine uplistings triggered comparable immediate increases. This suggests that genuineness of change influences which species attract attention more than how attention to a given species changes over time.

Our proxies for public and scientific interest—Wikipedia pageviews and Web of Science publications—carry some biases related to internet access, language, and media coverage. These measures may underrepresent regions with limited infrastructure and non-English research outputs. Nonetheless, they remain the most comprehensive global indicators available. Data limitations prevented inclusion of traits such as coloration, acoustic characteristics or deeper measures of cultural symbolism. In the quasi-experimental analysis, we cannot fully separate IUCN Red List category revisions from concurrent media or conservation campaigns and our estimates capture the combined informational effect. Temporal limitations also apply as yearly data may miss short-term spikes and the DiD model may not detect slow, long-term shifts.

In summary, public and scientific interest in birds is shaped primarily by charisma, familiarity, utilization, and Red List category. Yet Red List category changes produced only modest and asymmetric public responses, and the traits that predicted responsiveness to these changes differed sharply from the baseline drivers. Genuineness of change and increasing population trend, rather than any trait related to charisma, familiarity, or utilization, predicted the magnitude of public response. This disconnect suggests that the mechanisms generating baseline interest are not the same as those amplifying attention when a species’ Red List category changes. The disconnect between the predictors of baseline interest and those of change responsiveness suggests that simply updating a species’ Red List category is unlikely to shift public attention on its own. Proactive communication remains crucial, particularly for species that lack the charismatic or familiar traits that normally attract public interest ^5,13,56^. That even non-genuine uplistings triggered a significant immediate response further suggests an opportunity: communicating not only that a category has changed, but why, may help direct public attention where it is most warranted.

## Methods

### Data Acquisition

To investigate the drivers of attention, we compiled a global dataset for nearly all extant bird species. A complete description of all variables, their sources, and processing steps is provided in Supplementary methods and Table S9. Species taxonomy, scientific names, and synonyms followed the BirdLife International/HBW checklist version 10 (https://datazone.birdlife.org/about-our-science/taxonomy, accessed March 20th, 2026).

First, we collected data on a broad set of species-level traits. Phenotypic traits including body mass, relative beak and tail length was obtained from AVONET ^57^. Ecological traits included extent of occurrence, the latitudinal centroid of the species’ range, generation length, movement patterns, endemic status, utilization of anthropogenic habitat, and population trend. Cultural traits included indicators of a species’ relationship with humans, such as the consumptive use of species, number of years since described to science. These metrics were retrieved from the IUCN Red List database API (https://api.iucnredlist.org/). We also calculated the Web Capacity Index representing the internet usage within a species’ ranging countries, following Ladle et. al ^27^.

Second, we measured two indicators of attention. Scientific interest was quantified as the total number of articles in which a species’ scientific name appeared in the title, abstract or main text in the Web of Knowledge database via official API. This is a standard quantitative estimate of research effort toward individual species ^22,31,58^. Scientific names were used in preference to vernacular names, as binomial nomenclature avoids the ambiguity that can bias bibliometrics when common names refer to multiple entities (e.g., “teal” or “robin”) ^48^. Specifically, following the procedure a previous study utilised ^31^, we queried the Web of Knowledge database using topic searches (‘TS’). For each species, we used its scientific name along with synonyms as the search term, joined with the Boolean operator ‘OR’. This ensured that articles using alternate taxonomic names were not missed. We then recorded the total number of records published before 2026.

As Wikipedia data have been widely used to explore patterns of public interest in biodiversity, where total pageviews was often selected as a particularly useful metric ^31,59^, public interest was quantified as the total number of user pageviews for a species’ article across all available languages on Wikipedia. For each species, we first identified the corresponding Wikidata item identifier (QID) using the Wikidata Action API. We then verified each species’ QID and retrieved per-language page titles via the Wikidata API to query monthly user pageviews across all available Wikipedia language editions. Specifically, a species was considered to lack its own Wikipedia page when no dedicated species-level article existed (N = 238), even if it was mentioned within a broader article (e.g., a genus page) or treated as a subspecies under other species pages. As the Wikimedia pageviews API was introduced in August 2015 and has backfilled data to July 2015, we queried the API with the title and the language of each page to collect the monthly user pageviews (i.e., excluding views by bots) for the period between July 1, 2015, and December 31, 2025.

Third, to analyze the specific impact of species’ extinction risk category, we collected the complete assessment history for each species from the IUCN Red List of Threatened Species (Version 2025-2) ^60^. All Red List assessments for birds on the IUCN Red List are carried out by BirdLife International, as the designated Red List Authority for birds. For our analysis on the overall scientific and public interest, we selected the last assessment of each species and removed species classified as Data Deficient (DD, N = 36) because their extinction risk is too poorly known to classify, and removed Extinct (EX) and Extinct in the Wild (EW) species because we wanted to focus on extant species (N = 169). We then reclassified extinction risk into two classes to balance factor levels: Threatened, including Vulnerable (VU), Endangered (EN), and Critically Endangered (CR); Non-threatened, including Least Concern (LC) and Near Threatened (NT).

### Classification of IUCN Red List category changes

For our quasi-experimental analysis, we focused on Red List category change events occurring between 2019 and 2023 identifying 510 qualifying single-change events to form our “treated” group. As Wikipedia pageview data are available from 2017 to 2025, this time window ensures at least two years of observation for both pre-event and post-event baseline estimation. To allow for a detailed investigation into public response, these changes were categorized by direction (Uplist for an increase in extinction risk category; Downlist for a decrease) and a three-tiered measure of magnitude (Low, Moderate, or High).

We developed this magnitude classification to provide a more meaningful analysis than previous approaches. A simple numerical difference between categories, as used in some prior work ^53^, fails to distinguish between events of vastly different conservation significance. For example, a recovery from “Critically Endangered” to “Endangered” (−1 step) represents a greater reduction in extinction risk than a recovery from “Near Threatened” to “Least Concern” (−1 step). At the other extreme, modeling every unique category-to-category transition separately was statistically infeasible given our sample size, as it would have created many categories with too few observations, resulting in a critical loss of statistical power.

Our magnitude schema therefore aggregates these transitions into groups based on conservation significance. For uplistings, magnitude was defined by the destination category (Low: to NT; Moderate: to VU; High: to EN/CR), reflecting the importance of a species entering a higher extinction risk category. Conversely, for downlistings, magnitude was defined by the origin category (Low: from NT; Moderate: from VU; High: from EN/CR), reflecting the significance of a species recovering from a more severe level of threat.

To further examine whether the nature of the Red List category change and its communication shape public response, we incorporated two treatment-level variables. The genuineness of each category change indicates whether the category change reflected a real biological change (genuine) versus a revision due to improved knowledge, taxonomy, or methodology (non-genuine), which was provided by BirdLife International.

### Data analysis

We ran all analysis in R version 4.3.2 ^61^, using the *tidyverse* ^62^ and ‘*ggplot2*’ ^63^ suite for data handling and visualizations. We followed the general protocol for conducting and presenting results of regression-type analyses ^64^.

Our first analysis, modeling the drivers of baseline scientific and public interest, utilized Generalized Linear Mixed Models (GLMMs) fitted with the glmmTMB package ^65^. Our second analysis investigating the effect of IUCN Red List category changes employed a Difference-in-Differences (DiD) design. The causal effect was estimated using Linear Mixed-Effects Models (LMMs) with the lme4 package ^66^. This approach modeled the log-transformed pageviews as a function of the interaction between time relative to the event and treatment category. Finally, to investigate treatment effect heterogeneity, we conducted a two-stage meta-regression. In the first stage, we estimated the per-species effect size of the category change. In the second stage, we modeled these effect sizes using a Weighted Least Squares (WLS) linear model (lm function), with weights equal to the inverse of the variance of each effect size estimate.

For model validation, we used the suite of functions of the package ‘*performance*’ version 0.12.2 ^67^ to visually inspect model residuals and evaluate overdispersion, zero-inflation, and multicollinearity.

### Predictors of Baseline Public and Scientific Interest

To identify the primary drivers of baseline interest, we modeled both scientific interest (Web of Science article counts) and public interest (Wikipedia total pageviews) using Generalized Linear Mixed Models (GLMMs). To facilitate the interpretation of these results, we synthesized the variables into four thematic groups reflecting distinct drivers of human attention: Species charisma (mass, relative beak and tail length); Familiarity (years since described to science, web capacity index, anthropogenic habitat), Utilization (Use-Food, Use-Pet, Use-Sport, Use-Other), and Conservation-related attributes (IUCN Red List category, endemic status, extent of occurrence, latitude of range centroid, population trend). To account for phylogenetic non-independence, Order, Family, and Genus were included as random effects. Full model specifications and validation details are provided in the Supplementary Methods. We retained 10,509 bird species for analysis after removing the missing values. The final baseline interest model followed the formula (in R notation):

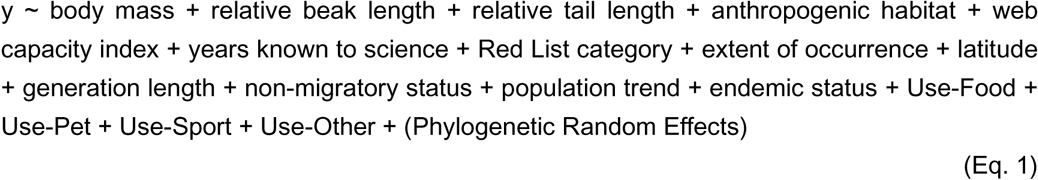

where y represents either the number of articles in the Web of Science (scientific interest) or the number of views on Wikipedia (public interest). The (Phylogenetic Random Effects) term was included as nested random intercepts to account for taxonomic non-independence, as species are not independent samples due to their shared evolutionary history. This mixed-effects structure is essential to avoid pseudo-replication and the potential for inflated statistical significance that arises from phylogenetically correlated data. For the public interest model, this included the full three-level structure of (1|Order) + (1|Family) + (1|Genus). For the scientific interest model, the (1|Order) term was excluded to resolve a singular fit identified during model diagnostics, resulting in the more parsimonious structure of (1|Family) + (1|Genus).

### Subset analyses on avian orders

To test the generality of our global models and explore potential clade-specific patterns, we repeated the baseline interest analyses on four major avian orders. These orders were chosen both for their large sample sizes, which support robust modeling, and for their distinct and contrasting ecological and cultural profiles. The selected orders were: Passeriformes, as the largest and most diverse avian order ^68^; Psittaciformes (parrots), a group defined by high visual charisma and its significant role in trade ^42^; Accipitriformes (most raptors), representing a classic form of charisma based on large size and predatory status ^69,70^; and Charadriiformes (shorebirds, gulls, and auks), representing an ecologically diverse but generally less charismatic clade.

The model structure for these subsets followed the same general formula as the main models (Eq. 1), with the (1|Order) random effect removed as it was invariant. Due to differences in the data distribution within each subset, minor adaptations to the models were sometimes necessary (e.g., adjusting the reference level for a categorical or resolving a singular fit). These subset-specific methodological details are fully documented in the Supplementary Methods.

### Analysis of Relative Interest

In addition to modeling absolute interest, we investigated the drivers of the relative balance between public and scientific interest. We first modeled the relationship between log-transformed public and scientific interest using a simple linear model and extracted the residuals for each species. Positive residuals identify species with greater public interest than predicted by their level of scientific interest, while negative residuals identify species with greater scientific interest than public interest. These residuals were then used as the response variable in a Gaussian Linear Mixed-Effects Model to identify the traits predicting this attention bias, following the same general formula as the simplified baseline model (Eq. 1). The (Phylogenetic Random Effects) term included the full structure of (1|Order) + (1|Family) + (1|Genus).

### Quasi-experimental analysis of IUCN Red List category change impacts

The preceding analysis provides a robust, static snapshot of the factors correlated with public interest across the bird species. It reveals what is popular, but it cannot explain how public interest responds to new information. The static IUCN Red List category used in that model, for example, is a powerful predictor, but it cannot tell us if the formal act of updating a species’ Red List category actually *causes* a significant shift in public interest. To address this critical question of causality, we designed a quasi-experimental analysis with a Difference-in-Differences (DiD) approach. By comparing the pageview trends of species that underwent a category change (the “treated” group) to those of a carefully matched control group that did not, the DiD design allows us to isolate the specific impact of the category update from broad, confounding background trends in public interest that affect all species over time. This design enables a stronger inference about the real-world impact of conservation assessments on public awareness.

The “treatment” in our design was an official IUCN Red List category change event. To ensure a robust analysis with a stable pre-event baseline and a sufficient post-event observation period, we restricted our sample to events occurring between 2019 and 2023, using pageview data from 2017 to 2025. Furthermore, to avoid the confounding effects of multiple interventions on the same species, we included only species that experienced a single Red List category change during this timeframe (N = 510, after excluding 197 species with more than one category changes). After excluding species with zero pageview variance across all years and those with missing data in matching covariates, our final treated group consisted of 479 qualifying category change events, comprising 354 uplistings and 125 downlistings.

To isolate the effect of the category change from general background trends, we constructed comparable control groups using a multi-stage matching procedure. For our primary analysis, we pooled all treated species (both uplists and downlists) and matched them to controls using 1:1 nearest neighbor matching. The distance between potential pairs was calculated based on a set of key covariates identified as strong predictors of public interest in our baseline models, ranked by effect size: log-transformed body mass, (signed) latitude of range centroid, web capacity index, years known to science, and binary indicators for Use-Food, Use-Pet, and Use-Sport. Furthermore, to ensure the validity of the comparison, we enforced exact matching on pre-change Red List category and genus (with family as a fallback). This hierarchical process prioritized controls that were taxonomically similar and had been recently reassessed without a category change.

For our focused sub-analyses, we created two distinct matched datasets. First, we isolated only the uplisted species and their potential controls and performed the matching procedure to create a dedicated uplist dataset. Second, we repeated this process for only the downlisted species to create a downlist dataset. This ensures that the comparisons within each sub-analysis are based on the most appropriate and balanced control group. Across all matching procedures, we successfully created control groups that were well-balanced with their respective treated groups. We also conducted an exploratory sensitivity analysis to determine whether a more direct measure of geographic proximity could improve matching quality and whether our substantive conclusions were robust to how geography was operationalized (see Supplementary Methods for details and full diagnostics).

We estimated the DiD effect using a series of Linear Mixed-Effects Models (LMMs) on log-transformed pageviews. For the primary analysis of all category changes, the model included interaction terms between time-since-event variables (‘post event’ for an immediate level change; ‘time since event’ for a sustained trend change) and a three-level factor for the treatment group (Control, Treated Uplist, Treated Downlist). This specification allowed us to simultaneously estimate the average impact of both uplist and downlist events relative to their matched controls. The model followed the general formula:

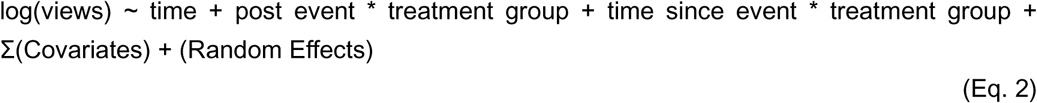

For the sub-analyses, we fitted separate models to the dedicated uplist and downlist matched datasets. These models replaced the ‘treatment group’ factor with a ‘change type’ factor to test for differences in effect based on the magnitude of the category change (Low, Moderate, High, and No Change controls). The general formula for these sub-models was:

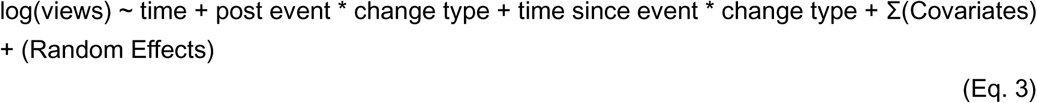

In all models, Σ(Covariates) represents the set of static species-level traits used as controls for any residual imbalance after matching. This set included the seven predictors identified as most influential in our baseline analysis: body mass, web capacity index, years known to science, latitude, Use-Food, Use-Pet, and Use-Sport. Additional traits predictive of public interest (generation length, extent of occurrence, relative beak and tail length, use-others, endemic status, anthropogenic habitat use, non-migrant status, and population trend) were also included in the full outcome model specification. The (Random Effects) represents a random intercept and random slope for each species (1 + time | scientific name) to account for individual variation in baseline popularity and trends.

To further analyze the roles of change direction and genuineness, we conducted a stratified DiD analysis that partitioned the matched pairs by whether the treated species underwent a genuine (N = 108) or non-genuine (N = 272) category change. The matching procedure and control assignments were held constant; only the treated pool was subset. This design ensures that any differences in estimates between strata reflect differences in the composition of the treated group rather than differences in matching quality or control selection. Within each stratum, we fitted a DiD model with the simplified covariate specification and random slopes as Eq.2.

After assessing the average treatment effect, we investigated which species traits predict a stronger or weaker public response to a Red List category change. This was achieved using a two-stage meta-regression analysis. In the first stage, we estimated the immediate effect size (the ‘post event’ coefficient) for each individual treated species using a linear model. In the second stage, these species-specific effect sizes were used as the response variable in a Weighted Least Squares (WLS) linear model, with species traits and the genuineness of change and whether the category change was mentioned in BirdLife International’s annual Red List news update (included as a control; see Supplementary Methods) as predictors, along with the inverse variance of the effect size estimate as weights. This analysis was performed on the full set of treated species, and then separately for the uplist and downlist subsets to test for differences in effect based on the magnitude of the Red List category change.

## Author Contributions

Haozhong Si: Conceptualization, Data curation, Formal analysis, Investigation, Methodology, Project administration, Resources, Software, Validation, Visualization, Writing - original draft, and Writing - review & editing.

Stuart H. M. Butchart: Conceptualization, Resources, Data curation, Methodology, Writing - review & editing.

Fan Yu: Conceptualization, Writing - review & editing. Changjian Fu: Conceptualization, Writing - review & editing.

Zhongqiu Li: Conceptualization, Funding acquisition, Project administration, Supervision, and Writing - review & editing.

Enrico Di Minin: Conceptualization, Funding acquisition, Project administration, Supervision, Writing - review & editing.

## Acknowledgments

We thank Josephine Burgin and Alex Berryman for their valuable help in extracting data of Red List category change history and verifying the genuineness of these changes. We would like to express our deepest gratitude to Chi Xu, and the open peer reviewers on Qeios, N. Riccardi, M. Lagisz, D. N. Jones, W. Fleming, J. C. Pena, and J. Guedes for comments on an earlier version of this manuscript. HS and EDM were funded by the European Union (ERC, BIOBANG, 101171602). Views and opinions expressed are however those of the authors only and do not necessarily reflect those of the European Union or the European Research Council Executive Agency. Neither the European Union nor the granting authority can be held responsible for them. Any opinions, findings, and conclusions or recommendations expressed in this material are those of the authors and do not necessarily reflect the views of the funding agencies.

