## Supplementary for "Global Scientific and Public Interest in Birds and their Conservation"

**This file includes:**

Supplementary Methods

Supplementary Results

Figure S1 to S7

Tables S1 to S9

**Supplementary Methods**

1. **Data acquisition and preparation**

**1.1 Compiling candidate factors**

A complete description of all variables, their sources, and processing steps is provided in Table S9. Species taxonomy, scientific names, and synonyms followed the BirdLife International/HBW checklist version 10 (<https://datazone.birdlife.org/about-our-science/taxonomy> , accessed March 20th, 2026). Where species were listed under different scientific names in earlier checklist versions (e.g., AVONET using HBW v5), we resolved name correspondences using shared BirdLife species identifiers (SISRecIDs). This identifier-based matching was used as the primary bridge for merging datasets compiled under different taxonomic versions, with scientific name matching applied as a secondary fallback.

Body mass, beak length (culmen), and tail length were obtained from AVONET (Tobias et al. 2022), a comprehensive dataset of functional traits for all bird species. AVONET follows HBW v5 taxonomy; records were matched to HBW v10 using SISRecIDs via the HBW v5 checklist, which carries both the v5 scientific name and the corresponding SISRecID for each species. Beak length and tail length were expressed relative to body mass by extracting residuals from phylogenetic regressions of each trait against log-transformed body mass, following (Santangeli et al. 2023). For species that were split between HBW v5 and v10, trait values from the v5 species were assigned to all resulting v10 species with the same SISRecIDs.

Extent of occurrence (EOO, km²), generation length (years), movement patterns, and population trend were retrieved from the IUCN Red List database via the official IUCN Red List API (https://api.iucnredlist.org, Version 4). To avoid ambiguity in interpreting migratory behaviors such as nomadism or altitudinal migration, we regrouped this into a binary non-migrant status: species classified as "Not a Migrant" were coded as non-migratory ‘Yes’; all other species with known movement patterns (Full Migrant, Altitudinal Migrant, Nomadic) were coded as ‘No’. Species with unknown or missing movement patterns (N = 33) were excluded. Endemic status was defined as species whose extant native range spans a single country, determined after resolving all IUCN sub-national and non-standard location codes to ISO 3166-1 alpha-3 country codes using a manually curated lookup table. A species was classified as using anthropogenic habitats if any IUCN habitat classification record for that species included urban, suburban, or artificial terrestrial environments (IUCN habitat classification scheme, version 3.1). The latitudinal centroid of each species' geographic range was derived from BirdLife International's global range maps

Consumptive use of species was coded as four binary variables: food (IUCN Use and Trade code 1), pet (code 13), game (code 15), and other use, retrieved from IUCN Red List Database (IUCN general use and trade classification scheme, version 1.0). Years known to science was calculated as the difference between the current year (2026) and the year of taxonomic description, extracted from the authority string in the IUCN taxon record. We also collected the ‘Population trend’ status (Increasing, Stable, Decreasing, Unknown) from the IUCN Red List Database.

The Web Capacity Index (WCI) represents the proportion of global internet users residing in countries where a species has an extant native range, providing a measure of the potential human digital audience for each species. Country-level internet penetration rates (percentage of individuals using the internet) were obtained from the International Telecommunication Union (ITU) ICT indicators database (<https://data.worldbank.org/indicator/IT.NET.USER.ZS> , accessed March 27^th^, 2026) for the reference year 2019, and national population counts were obtained from the World Bank (<https://data.worldbank.org/indicator/SP.POP.TOTL> , accessed March 27^th^, 2026) for the same year. Absolute internet user counts per country were computed as the product of the penetration rate and population, then summed across all range countries and divided by the global total.

Missing ITU penetration values for 2019 were imputed sequentially using three methods: (1) linear interpolation within each country's time series between the nearest available preceding and following years, which is defensible given the near-monotonic trajectory of internet adoption; (2) the median penetration rate of the country's UN sub-region for 2019, applied to countries with no data in adjacent years. We selected 2019 as the reference year to avoid anomalous patterns in both Wikipedia traffic and online research activity associated with the COVID-19 pandemic (2020–2021), while remaining representative of the digital landscape during our Wikipedia pageview observation period.

Additionally, the IUCN Red List is a cornerstone of global conservation policy (Rodrigues et al. 2006; Betts et al. 2020), and Red List category was suggested to be a significant driver of scientific and public interest (Roll et al. 2016; Ladle et al. 2019; Mammola et al. 2023). For our analysis on the overall scientific and public interest, we collected the most recent assessment of each species and removed species with a Data Deficient (DD, N = 36), Extinct (EX) or Extinct in the Wild status (EW) (N = 169) because the conservation risk of such species is ambiguous and may bring addition bias to our analyses. We then reclassified Red List categories into two categories to balance factor levels: Threatened (including ‘Vulnerable’, ‘Endangered’, and ‘Critically Endangered’), Non-threatened (including ‘Least Concern’ and ‘Near Threatened’). These assessments reflect the most recent global evaluations as of 2024 and were treated as static values. For metrics retrieved from IUCN Red List Database, we used the Python package ‘*requests*’ version 2.32.3 (Reitz 2024) to query the IUCN Red List API v4 and access the IUCN Red List of Threatened Species version 2025(2).

Specifically, to analyze the effect of Red List category changes, we collected a complete history of IUCN Red List assessments for all bird species. We then identified all instances where a species' assessment was updated, noting whether this resulted in a Red List category change. To ensure a robust analysis with a sufficient pre-event baseline and post-event observation period, we restricted our study to events occurring between 2019 and 2023, using pageview data from 2017 to 2025. Furthermore, to avoid confounding effects from multiple interventions, only species that experienced a single category change during this timeframe were included in the treated group. This filtering process yielded a total of 510 qualifying category change events. We also classified the category changes by direction (Uplist for an increase in threat level; Downlist for a decrease). For our sub-analyses, we further classified category changes by their magnitude, where Uplist magnitude was defined by the destination category (Low: to NT; Moderate: to VU; High: to EN/CR), while Downlist magnitude was defined by the origin category (Low: from NT; Moderate: from VU; High: from EN/CR). This process allowed for a detailed investigation into how different types and magnitude of Red List category changes impact public attention.

To further examine whether the nature of the Red List category change and its communication shape public response, we incorporated two treatment-level variables. The genuineness of each category change indicates whether the IUCN category change reflected a real biological change (genuine) versus a revision due to improved knowledge, taxonomy, or methodology (non-genuine), which was provided by BirdLife International. Whether the category change was featured in BirdLife's annual Red List update news was manually collected from the BirdLife International websites (including <https://datazone.birdlife.org/articles> and <https://www.birdlife.org/news>) and verified by the authors.

1.2 **Measures of scientific and public interest**

We gathered data on two indicators reflecting attention towards bird species, related to scientific and public interests.

We measured scientific interest as the total number of peer-reviewed articles and reviews in which a species' name appeared in the title, abstract, or keywords. This is a standard quantitative estimate of research effort toward individual species (dos Santos et al. 2020; Adamo et al. 2021; Mammola et al. 2023). Scientific names were used in preference to vernacular names, as binomial nomenclature avoids the ambiguity that can bias bibliometrics when common names refer to multiple entities (e.g., "teal" or "robin") {Correia, 2017 #282}. We retrieved this data via the Clarivate Web of Science Expanded API (https://developer.clarivate.com/apis/wos) using topic search (TS=) queries. For each species, the accepted scientific name and all recognized synonyms from the BirdLife Datazone were joined with the Boolean operator ‘OR’ as the search term. We queried the Web of Knowledge all-databases endpoint (databaseId=WOK), which returns a server-side deduplicated count across multiple databases. This broader union was preferred over the WOS Core Collection alone, as the Core Collection alone produced zero counts for 34% of species compared to 6% for the all-databases union, likely with substantial research on tropical and specialist taxa being published in regional and specialist journals indexed outside the Core Collection. We then recorded the total number of records published before 2026.

We measured public interest for each species as the total number of pageviews across the languages where the species is represented on Wikipedia. As one of the top 10 most visited websites in the world (<https://www.similarweb.com/top-websites>, accessed on Sep 1, 2024), Wikipedia provides a vast source of information for bird enthusiasts, with the majority of bird species containing a page in this digital encyclopedia. Wikipedia data has also been widely used to explore patterns of public interest in biodiversity, where total pageviews was often selected as a particularly useful metric (Vardi et al. 2021; Mammola et al. 2023).

To accurately retrieve pageview statistics for the world's bird species, each taxon must first be mapped to its corresponding Wikidata item identifier (QID). Wikidata is a structured, linked-data knowledge base that underpins Wikipedia's infoboxes and interlanguage links, and each Wikipedia article is associated with exactly one Wikidata item. We developed a sequential, multi-round pipeline to assign a Wikidata ID to each bird species recognized in our taxonomic backbone. The pipeline proceeds from a taxonomic identifier through progressively broader search strategies, followed by automated validation and manual review. All data retrieval was conducted between 30-03-2026 to 14-05-2026.

We mapped each bird species (n = 11,185 species) to its corresponding Wikidata item identifier (QID) through a four-round sequential pipeline. Round 1 translated the IUCN Species Information Service identifier (SISID, share identical values with SISRecID) to an Open Tree of Life identifier via the OneZoom API (<https://www.onezoom.org> ), then retrieved the linked Wikidata ID from OneZoom's cross-referenced identifier mappings. Round 2 queried the Wikidata SPARQL endpoint with the scientific name, matching against property P225 (taxon name) to obtain the QID directly. Round 3 processed species still unmatched after the first two rounds—as well as species whose cumulative Wikipedia pageviews fell below 10,000—through the English Wikipedia Search API, filtering results to exclude list articles, higher-taxon pages, and non-species entries, then extracting the Wikidata ID from the matched page. Round 4 comprised manual curation of any remaining unmatched taxa.

We validated the resulting mapping in two stages. First, each QID was checked against Wikidata (property P31 "instance of" for valid taxon types; property P105 "taxon rank" restricted to species) to confirm it referred to a bird species. Second, duplicate QIDs, arising from taxonomic disagreements between HBW-BirdLife and Wikipedia, were resolved by setting the QID to missing where the relevant Wikipedia article treated the taxon as a subspecies without a dedicated page. The final mapping contained 10,947 entries, representing 97.9% of the initial taxonomic backbone. Species with no dedicated species-level article existed was considered to lack its own Wikipedia page (N = 238), even if it was mentioned within a broader article (e.g., a genus page) or treated as a subspecies under other species pages. We then used species’ QID to retrieve per-language page titles via the Wikidata API and query monthly user pageviews across all available Wikipedia language editions. As the Wikimedia pageviews API was introduced in August 2015 and has backfilled data to July 2015, we queried the API with the title and the language of each page to collect the monthly user pageviews (i.e., excluding views by bots) for the period between July 1, 2015, and December 31, 2025.

**1.3 Data exploration**

Data exploration was conducted following the protocol described by Zuur, Ieno, and Elphick (Zuur et al. 2010). Prior to model construction, we visually inspected variable distributions, the presence of outliers, multicollinearity among predictors, and the balance of factor levels.

We log-transformed body mass and extent of occurrence to homogenize their distributions and minimize the effect of outliers. All continuous variables were scaled (to a mean of zero and a standard deviation of one) to obtain comparable effect sizes and facilitate the convergence of regression models. Multicollinearity testing with Pearson’s r correlations indicated no problematic multicollinearity (all |r| < 0.60), and all Variance inflation factors (VIFs) fell below 2.0.

Upon inspecting the distribution of the number of articles in the Web of Science (scientific interest) and the number of views on Wikipedia (public interest), we identified several outliers. We excluded the Greater Honeyguide (*Indicator indicator*), Melodious Blackbird (*Dives dives*), Whooper Swan (*Cygnus cygnus*) and Socotra warbler (*Incana incana*) due to the generation of numerous unrelated results from topic searches of their scientific names.

2. **Baseline interest model**

**2.1 Fitting and validation of the interest baseline models**

To identify the primary drivers of baseline interest, we modeled two response variables: scientific interest (Web of Knowledge article counts) and public interest (total Wikipedia pageviews). Given that both are count data, we used Generalized Linear Mixed Models (GLMMs) assuming a Poisson error structure and a log-link function using *glmmTMB* package (Brooks et al. 2017). We fitted the baseline models for both scientific and public interest, which are the ones interpreted and visualized in the main manuscript. They were specified as Generalized Linear Mixed Models (GLMMs) using the *glmmTMB* package in R (Brooks et al. 2017), with a negative binomial error distribution for public interest model, and zero-inflated truncated negative binomial error distribution for scientific interest model. The general formula for the final models was (in R notation):

y ~ body mass + relative beak length + relative tail length + anthropogenic habitat + web capacity index + years known to science + Red List category + extent of occurrence + latitude + generation length + non-migratory status + population trend + endemic status + Use-Food + Use-Pet + Use-Sport + Use-Other + (1|Order) + (1|Family) + (1|Genus)

(Eq. S2)

where y represents either the number of articles in the Web of Science (scientific interest) or the number of views on Wikipedia (public interest). Note that the latitude refers to the absolute latitude of range centroid. We included Order (factor with 36 levels), Family (246 levels), and Genus (2353 levels) as random intercept factors to account for taxonomic non-independence of samples. The model for scientific interest, following the singularity resolution described previously, used a simplified random effects structure where the (1|Order) term was removed.

We conducted a thorough validation of both final models using the *performance* package (Lüdecke et al. 2021). Diagnostic checks confirmed that there were no significant issues with overdispersion, zero-inflation, or multicollinearity (all Variance Inflation Factors < 3). Besides, our simplified models demonstrated strong explanatory power, with the selected fixed effects accounting for a substantial portion of the variance in both public interest (Marginal R² = 0.585) and scientific interest (Marginal R² = 0.609).

**2.3 Analysis of relative interest**

In addition to modeling absolute public and scientific interest, we conducted an analysis to understand the drivers of relative interest between the two domains.

First, to quantify the balance of interest for each species, we modeled the relationship between log-transformed public interest and log-transformed scientific interest using a simple linear model (lm function in R). We then extracted residuals for each species from this regression. In this framework, positive residuals identify species with greater public interest than predicted by their level of scientific interest, while negative residuals identify species with greater scientific interest than public interest.

Second, to identify the traits associated with this bias in interest, we used these residuals as a continuous response variable in a Gaussian Linear Mixed-Model. This model followed the same general formula as the simplified baseline model (Eq. S2), allowing us to evaluate which traits predict a species' deviation from the expected public-scientific interest relationship.

**2.4 Subset analyses**

To test the generality of our global models and explore potential clade-specific patterns, we repeated the baseline interest analysis on four major bird orders. These orders were chosen both for their large sample sizes, which support robust modeling, and for their distinct and contrasting ecological and cultural profiles. The selected orders were: Passeriformes, as the largest and most diverse bird order (N = 6,315) (Ericson et al. 2003); Psittaciformes (parrots, N = 384), a group defined by high visual charisma and its significant role in trade (Frynta et al. 2010); Accipitriformes (most raptors, N = 248), representing a classic form of charisma based on large size and predatory status (Buechley et al. 2019; Horgan et al. 2021); and Charadriiformes (shorebirds, gulls, and auks, N = 366), representing an ecologically diverse but generally less charismatic clade.

The model structure for each subset analysis followed the same general formula and predictor set as the main models (Eq. 1), though several necessary, subset-specific adaptations were made to ensure model stability and validity. For all four subsets, the (1|Order) random effect was removed as it was invariant. Additionally, the public interest models for Passeriformes and Charadriiformes, along with the scientific interest models for Accipitriformes and Psittaciformes initially produced singular fits with a (1|Family) + (1|Genus) structure. Following best practice, this was resolved by simplifying the structure to (1|Genus). All such adaptations are reflected in the full model output tables (Table S2).

**3.1 Quasi-experimental analysis of Red List category change impacts**

To assess the causal impact of Red List category changes on public interest, we employed a quasi-experimental Difference-in-Differences (DiD) design. This approach is superior to simple correlational analyses as it allows for the isolation of the treatment effect from confounding trends that may affect public interest in all species over time (Abadie 2005). The "treatment" was defined as an official update to a species' IUCN Red List category. The analysis was designed to evaluate two metrics of public response: (1) the immediate effect, i.e., an instantaneous level change in pageviews in the year of the update, and (2) the sustained effect, i.e., a change in the long-term slope of the pageview trend following the update.

**3.1 Data filtering and event definition**

To construct a robust analytical sample, we applied a multi-stage filtering protocol to the complete IUCN assessment history for all bird species, using Wikipedia pageview data from January 2017 to December 2025.

First, we restricted our analysis to category change with the publish year between 2019 and 2023. This time window was chosen to ensure that every event in our sample had both a stable pre-event baseline period of at least two years (2017-2018) and a post-event observation period of at least two years (2024-2025). This allows for a reliable estimation of both pre-event and post-event trends.

Second, to avoid the complex confounding effects of various interventions, we only included species that experienced exactly one Red List category change within our full data window (2017-2025). Any species that was updated more than once during this period was excluded from the analysis (N = 197). This ensures that the pre-event baseline for any given event is not affected by the previous, recent category change.

Third, we excluded species without matched Wikidata ID (N = 234), which have zero pageview variance across the entire study and provide no measurable signal of public interest. Species with missing data in any matching covariate (N = 729) were also excluded via complete-case filtering. Finally, species with "Unknown" population trend (N = 16) were excluded to avoid ambiguity in this key confounding variable. Additionally, we regrouped the raw movement pattern variable into a three-level non-migrant status (Yes/No/Unknown), following the classification used in prior analyses within this study. Species with unknown non-migrant status (N = 2) were excluded.

After applying this filtering protocol, our final "treated" group consisted of 479 unique category change events. To allow for a detailed investigation into public response, these changes were categorized by direction (Uplist for an increase in threat level, N = 356; Downlist for a decrease, N = 125) and a three-tiered measure of magnitude (Low, Moderate, or High).

We developed this magnitude classification to provide a more meaningful analysis than previous approaches. A simple numerical difference between categories, as used in some prior work (Van Huynh 2023), fails to distinguish between events of vastly different conservation significance. For example, a recovery from “Critically Endangered” to “Endangered” (-1 step) and a recovery from “Near Threatened” to “Least Concern” (-1 step) represent fundamentally different conservation outcomes. On the other extreme, modeling every unique category-to-category transition separately was statistically infeasible given our sample size, as it would create many categories with too few observations, resulting in a critical loss of statistical power.

Our magnitude schema therefore aggregates these transitions into theoretically coherent groups based on conservation significance. For uplist changes, magnitude was defined by the destination category (Low: to NT; Moderate: to VU; High: to EN/CR), reflecting the importance of a species entering a higher threat level. Conversely, for downlist changes, magnitude was defined by the origin category (Low: from NT; Moderate: from VU; High: from EN/CR), reflecting the significance of a species recovering from a more severe level of threat.

**3.2 Matched control group construction and validation**

To validly estimate the causal effect of the Red List category change, a comparable control group was constructed for each analysis using a multi-stage matching procedure with the *MatchIt* package in R (Ho et al. 2011). This procedure was designed to create a control group that was as similar as possible to the treated group on a wide range of observable covariates, thereby reducing bias from confounding variables.

We employed a hierarchical 1:1 nearest neighbor matching algorithm without replacement. The matching was performed sequentially to prioritize the highest-quality controls. First, we defined two tiers of potential controls: a "Gold Standard" pool of species that were reassessed by the IUCN in the same year as a treated species but whose Red List category was reaffirmed, and a "Silver Standard" pool of species that were not assessed during the event window. The matching algorithm then proceeded in four stages for each event year: (1) it first attempted to find a Gold Standard control within the same Genus; (2) failing that, within the same Family; (3) failing that, it searched for a Silver Standard control within the same Genus; and finally, (4) within the same Family. This hierarchical approach ensured that the closest possible phylogenetic and temporal matches were prioritized.

The distance between potential pairs was calculated based on a set of key covariates identified as strong predictors of public interest in our baseline models, ranked by effect size: log-transformed body mass, (signed) latitude of range centroid, web capacity index, years known to science, and binary indicators for Use-Food, Use-Pet, and Use-Sport. Furthermore, to ensure the validity of the comparison, we enforced exact matching on pre-change IUCN Red List category and genus (with family as a fallback). For the uplist and downlist sub-analyses, we additionally included population trend as an exact matching criterion, given its mechanistic link to Red List category assessment (e.g., declining species are far more likely to be uplisted).

This entire matching procedure was conducted independently for our three main analyses. A primary matching process was run on the full set of 479 treated events to create an overall matched dataset (380 successfully matched pairs, max |SMD| = 0.141). Subsequently, two separate matching processes were run exclusively on the uplist and downlist subsets using the stricter matching criteria to generate dedicated, optimally balanced datasets for each sub-analysis (Uplist: 130 pairs, max |SMD| = 0.191; Downlist: 192 pairs, max |SMD| = 0.131).

The validity of the matching was confirmed by assessing covariate balance before and after each procedure. We calculated the Standardized Mean Difference (SMD) for all covariates, with a target threshold of |SMD| < 0.20 indicating a well-balanced sample. As detailed in the covariate balance tables (Table S8), the matching procedures were highly successful, substantially reducing the SMD for all covariates and creating valid control groups for all three analyses. The uplist sub-analysis showed a minor residual imbalance in the Use-Sport variable (SMD = 0.191), attributable to the inherently limited pool of control species with declining population trends. Any minor residual imbalances were statistically controlled by including the respective covariates as fixed effects in the final regression models.

**3.3 Primary DiD model (overall effect)**

We used a series of Linear Mixed-Effects Models (LMMs), fitted with the *lme4* package in R (Bates et al. 2015), to estimate the DiD effect of Red List category changes on log-transformed Wikipedia pageviews. All models included a standard set of random effects to account for individual variation (1 + (time | scientific name)).

To assess the average impact of all category changes, we fitted a primary DiD model to the overall matched dataset. The model's key predictors were the interaction terms between ‘post event’ or ‘time since event’ variables and a three-level treatment group factor. This factor was coded as Control (reference, no Red List category change), Treated Uplist, and Treated Downlist. The model followed the general formula:

log(views) ~ time + post event * treatment group + time since event * treatment group + Σ(Covariates) + (Random Effects)

(Eq. S3)

In this specification, the ‘post event: treatment group’ interaction term estimates the average immediate effect (level change), while the ‘time since event: treatment group’ interaction term estimates the average sustained effect (slope change) for uplist and downlist events, each relative to the control group. The covariates included the seven matching distance variables (log-transformed body mass, web capacity index, absolute latitude of range centroid, years known to science, Use-Food, Use-Pet, Use-Sport) along with population trend, generation length, extent of occurrence, relative beak and tail length, Use-Others, endemic status, anthropogenic habitat use, and non-migratory status, i.e., the full set of predictors for public interest in our baseline model. We note that latitude enters the models in two forms: signed latitude is used as a matching distance variable (preserving north-south hemisphere discrimination for control selection), while absolute latitude is used in all outcome models (reflecting the conceptual expectation that distance from the equator, rather than hemisphere per se, drives variation in public attention). The model converged cleanly with the ‘bobyqa’ optimizer and showed no rank deficiency or singularity warnings

.

**3.4 Sub-analysis DiD models**

For the sub-analyses, we fitted separate models to both the uplist and downlist matched datasets. These models replaced the ‘treatment group’ factor with a ‘change type’ factor to test for differences in effect based on the magnitude of the Red List category change (Low, Moderate, High, and No Change controls). This allowed for a direct estimation of the immediate and sustained effects for each magnitude level, relative to the matched control group within each subset. The random effects and covariates stayed the same as in Eq. S3. The general formula for these sub-models was:

log(views) ~ time + post event * change type + time since event * change type + Σ(Covariates) + (Random Effects)

(Eq. S4)

To further analyze the roles of change direction and genuineness, we conducted a stratified DiD analysis that partitioned the matched pairs by whether the treated species underwent a genuine (N = 108) or non-genuine (N = 272) category change. The matching procedure and control assignments were held constant; only the treated pool was subset. This design ensures that any differences in estimates between strata reflect differences in the composition of the treated group rather than differences in matching quality or control selection. Within each stratum, we fitted a DiD model with the simplified covariate specification and random slopes as Eq.S3.

**3.5 Meta-regression analysis on factors of treatment effect**

Finally, to identify species-level traits that predict sensitivity to a Red List category change, we conducted a two-stage meta-regression analysis.

In addition to the full set of static species-level traits from our baseline analysis, we considered two treatment-level characteristics that may moderate the effect size, whether the category change was genuine (‘genuine_change’) and, whether it was mentioned in BirdLife International's annual news update (‘mentioned_BL_annual’). This provides a direct test of whether formal conservation communication amplifies public attention. Both variables are binary and were treated as factors in the analysis.

First, we estimated the immediate effect size for each species by fitting a simple linear model (lm(log(views) ~ time + post event)) for each of the 380 treated species individually. From each model, we extracted the coefficient for ‘post event’ as the species' effect size and its corresponding standard error.

In the second stage, we used these effect size estimates as the response variable in a Weighted Least Squares (WLS) linear model. The model included the full set of static species traits (except for the static Red List category) from our baseline analysis as predictors, along with ‘genuine_change’ and ‘mentioned_BL_annual’. The model included a comprehensive set of predictors drawn from our baseline analysis, following the formula:

Effect size ~ body mass + latitude + range size + web capacity index + years known to science + generation length + relative beak length + relative tail length + Use-Food + Use-Pet + Use-Sport + Use-Others + endemic species + anthropogenic habitats + non-migratory status + population trends + genuine change + mentioned in annual news (+ change direction/magnitude)

(Eq. S5)

To give more influence to more precisely estimated effects, the model was weighted by the inverse of the variance of each effect size estimate (weights = 1 / (std.error^2)). This analysis was performed first on the full set of treated species (N = 380), and then separately on the uplist (N = 130) and downlist (N = 192) subsets to evaluate predictive patterns for different direction of the category change. As only 51 species have been mentioned during the period, we also fitted a simpler specification excluding the mentioned_BL_annual variable to assess its influence on model fit.

The validity of each meta-regression model was assessed using the overall F-statistic, which tests whether the model as a whole explains a significant portion of the variance in effect sizes, and the adjusted R-squared (Adj. R²) value, which quantifies the model's explanatory power. Diagnostic plots of the weighted residuals were inspected to check for patterns indicating violations of model assumptions. As reported in the main text, the full meta-regression model pooling all treated species was statistically significant (F_(20, 359)_ = 2.13, p = 0.004, Adj. R² = 0.056), with the genuineness of change emerging as the strongest individual predictor. The simpler specification excluding mentioned_BL_annual showed a similar fit (F_(19, 360)_ = 1.94, p = 0.011, Adj. R² = 0.045). The uplist and downlist subset models were not significant at the α = 0.05 level. The modest Adj. R² values (0.01–0.06) are consistent with the inherent difficulty of predicting treatment effect heterogeneity from noisy per-species estimates, and the significant F-tests confirm that the models capture systematic variation beyond chance.

**3.6 Genuine change analysis**

As introduced in Section 3.5, "genuine" Red List category changes reflect a real improvement or deterioration in a species' conservation status, and "non-genuine" changes, which result from taxonomic revisions, improved knowledge, or changes in assessment methodology. To complement the meta-regression analysis, which tested whether change authenticity predicts between-species effect sizes, we conducted a series of interaction models to directly test whether the within-species public response over time differs by the genuineness of change.

Among the 479 treated species, 135 had genuine changes and 344 had non-genuine changes. Genuine changes were substantially more common among uplistings (96 of 154 total uplist treated species) than among downlistings (39 of 356).

We fitted three complementary model specifications to assess the effect of the genuineness of change. First, a treated-only LMM was fitted to the full set of treated species from the overall matched dataset (N = 380). This model tested whether, among species that underwent a category change, the immediate and sustained effects differed by the genuineness of change. The model followed the formula:

log(views) ~ time + Σ(Covariates) + post event * genuine change + time since event * genuine change + (1 + time | scientific name)

(Eq. S6)

Second, to provide a more stringent causal test, we extended the primary DiD model (Eq. S3) to include a three-way interaction with change authenticity, using the full matched dataset including controls (N = 742 species, 6,678 observations):

log(views) ~ time + Σ(Covariates) + post event * treatment group * genuine change + time since event * treatment group * genuine change + (1 + time | scientific name)

(Eq. S7)

In this specification, the three-way interaction terms (e.g., 'post event: treatment group [Treated Downlist]: genuine change') estimate whether the DiD effect, already differenced by treatment status, further differs by the genuineness of change. This model was fitted only when each subgroup contained at least 20 species. Controls, which by definition have no category change, were assigned as non-genuine. The model converged cleanly with the ‘bobyqa’ optimizer and showed no rank deficiency.

Third, we repeated the treated-only analysis (Eq. S6) separately for the uplist-only matched dataset (N = 130 treated species, 82 genuine) and the downlist-only matched dataset (N = 192 treated species, 13 genuine) to assess whether the role of change authenticity differed by direction.

**3.7 News mention analysis**

As introduced in Section 3.5, we investigated whether formal conservation communication amplifies the public impact of Red List category changes by testing whether species mentioned in BirdLife International's Red List update news showed a stronger pageview response. To complement the meta-regression and provide a within-species temporal test, we fitted a series of interaction models analogous to those in Section 3.6. A species was coded as mentioned (mentioned_BL_annual = 1) if its category change was featured in the annual update for its event year, and 0 otherwise.

Among the 510 treated species in the full (pre-filtering) sample, only 51 were mentioned, 34 uplisted and 17 downlisted species. Consequently, analyses involving this variable should be interpreted as exploratory due to limited statistical power.

Unlike the genuine change analysis, a full-DiD or triple-difference extension is not applicable here, because control species did not undergo a Red List category change and therefore cannot be "mentioned in the annual news for a category change." We therefore restricted our analysis to a treated-only design. A treated-only LMM was fitted to the full set of treated species from the overall matched dataset (N = 380 treated species, 41 mentioned species matched):

log(views) ~ time + Σ(Covariates) + post event * mentioned_BL_annual + time since event * mentioned_BL_annual + (1 + time | scientific name)

(Eq. S8)

The parameters 'post event: mentioned_BL_annual' and 'mentioned_BL_annual: time since event' estimate the additional immediate and sustained effect associated with being featured in the annual news update. However, the post-event × news mention interaction term was dropped from models due to rank deficiency, as species mentioned all went through Red List category changes at the same year.

As with the genuine change analysis, we repeated this specification separately for the uplist-only and downlist-only matched datasets to explore direction-specific patterns. The uplist subset retained 32 of the 35 mentioned uplist species, yielding a reasonably powered comparison. However, the downlist subset retained only 2 of the 16 mentioned downlist species. These mentioned downlistings tend to be recoveries with rare combinations of pre-change IUCN category and population trend, making them difficult to match under the exact matching criteria. Given this, the downlist-only news mention model could not be reliably estimated and is not reported.

As noted in Section 3.5, both genuine_change and mentioned_BL_annual were also included as predictors in the meta-regression analysis. The interaction-based analyses presented here and in Section 3.6 test whether these variables moderate the within-species treatment effect over time, whereas the meta-regression tests the complementary question of whether they predict the magnitude of the between-species effect size. Results from both analytical frameworks are reported jointly in the main text.

**3.8 Geographic proximity matching as an exploratory sensitivity analysis**

The matching distance formula in our main analysis (Section 3.2) incorporates two geographic proxies, latitude of range centroid and web capacity index (WCI). Latitude approximates broad biogeographic position, while WCI captures country-level internet infrastructure and indirectly reflects the geographic distribution of Wikipedia readership. Together, these variables serve as parsimonious geographic controls. However, neither directly measures the spatial overlap between a species' actual geographic range and the regions from which Wikipedia pageviews originate. We therefore conducted an exploratory sensitivity analysis to determine whether a more direct measure of geographic proximity could improve matching quality and whether our substantive conclusions were robust to how geography was operationalized.

A direct approach would compute the pairwise spatial overlap of species' continuous range polygons. We rejected this option on both practical and conceptual grounds. Practically, computing pairwise polygon intersections for over 10,000 species is computationally prohibitive. Conceptually, even if feasible, spatial adjacency does not equate to shared public attention. Two neighbouring countries may share a border and similar avifauna, yet diverge substantially in the size of their online readership, the institutional capacity of their conservation organisations, and the level of public engagement with biodiversity information, all of which affect Wikipedia pageview patterns. A continuous range overlap metric would falsely equate a species spanning, for example, Singapore and Malaysia with a species spanning two European countries with comparable digital infrastructures, despite the former pair likely exhibiting far greater asymmetry in public attention.

We therefore adopted a discrete, tractable alternative: country presence data, extracted from the IUCN range database. For each species, we constructed a binary vector indicating presence or absence in each of 250 countries, ignoring seasonality and species origin. From this species × country matrix, we computed pairwise Jaccard distance as:

d_Jaccard(A, B) = 1 − |countries(A) ∩ countries(B)| / |countries(A) ∪ countries(B)|

where a distance of 0 indicates identical country sets and a distance of 1 indicates no shared countries. Country presence data were available for 10,994 bird species, including all treated species in our sample.

To integrate geographic and trait-based information, we first tested an α-weighted linear combination: d_combined = α × d_Jaccard + (1 − α) × d_traits, where d_traits is the standardized Euclidean distance computed from the five non-geographic matching covariates (body mass, years known to science, Use-Food, Use-Pet, and Use-Sport). We tested α values ranging from 0.3 to 0.95. This approach proved entirely ineffective: all α values produced identical matched pairs. The exact matching constraints (loosen to b_status + genus/family/order) partition the species pool into small, homogeneous strata, and within any given stratum the α-weighted distance formula had no discriminating power. This suggests that the match was determined by whichever single control happened to be available, regardless of the weighting scheme. This finding revealed a fundamental tension between exact matching, which enforces strict comparability on key confounders, and distance-based matching, which requires a meaningful gradient of similarity within those constraints to differentiate among candidate controls.

Recognising this limitation, we reframed the approach lexicographically: rather than weighting geography and traits simultaneously, we prioritised geographic proximity as a first-stage criterion and fell back to trait-based matching only when no geographic peer was available. Specifically, within each exact match stratum, the algorithm first attempted to match each treated species to a control with Jaccard distance less than 1 (i.e., at least one shared country), selecting the closest match by trait distance when multiple candidates existed. For treated species with no such geographic peer, it then matched using trait distance alone, ensuring that the stringent geographic requirement did not come at the cost of unmatched treated units. This lexicographic design reflects the correct decision hierarchy: geographic proximity is a desirable matching criterion when feasible, but should not force the exclusion of species for which no suitable geographic control exists.

Under this lexicographic approach, 37% of matched pairs were assigned via the geographic-first pathway, with a median Jaccard distance of 0.667 among these pairs, indicating meaningful country overlap. The remaining 63% of pairs had zero country overlap within their exact match strata and were matched using trait distance alone. This result is structural rather than algorithmic, and the underlying country presence data reveal its origin: a majority of treated species (57.5%) are endemic to a single country. For a single-country endemic to find a geographic peer, the exact match stratum, defined by Red List category and taxonomic group, must contain a control species that shares that specific country. When the control pool within that stratum consists almost entirely of species from different continents, no geographic match is possible under any matching rule. This constraint is compounded by the geographic concentration of treated species in centres of avian endemism, Indonesia, Colombia, Brazil, Peru, and Ecuador collectively account for over half of all treated species, where many species have ranges confined within a single national border. A passerine endemic to the Peruvian Andes and a passerine endemic to the Indonesian archipelago, for example, may belong to the same Order and Red List category yet share no countries, and the country-presence metric provides no basis for distinguishing between them on geographic grounds.

The lexicographic geographic matching produced a larger sample (466 pairs, 96.3% match rate) but at the cost of worse covariate balance (max |SMD| = 0.236 vs. 0.141 in the main analysis). The DiD estimates were qualitatively consistent with the main results. The uplist immediate effect remained significant and of comparable magnitude (0.079, p = 0.009 vs. main: 0.064, p = 0.029), and the downlist and news mention effects remained non-significant, indicating that the substantive findings are not artefacts of the specific geographic specification.

This sensitivity analysis demonstrates that our main approach of using latitude centroid and WCI as geographic proxies is not merely a simplification but a deliberate methodological choice: these continuous variables are sufficiently coarse to accommodate the global nature of Wikipedia readership while sufficiently informative to control for broad geographic confounding. Country-level presence data, by contrast, imposes a binary filter that is too strict, it penalises species pairs whose ranges are geographically adjacent but politically separated, and it cannot capture the continuous gradient of public attention across space. Second, it provides reassurance that our core substantive conclusions are robust to how geographic information is incorporated into the matching design. The geo-matching experiment confirms that we have considered, tested, and, on explicit methodological grounds, not adopted a more elaborate geographic proximity measure, thereby strengthening the transparency and defensibility of our final modelling choices.

**3.9 Simplified covariate specification as a robustness check**

The primary outcome models (Eqs. S3–S8) include the full set of baseline species traits, i.e., 15 fixed effects beyond the DiD interaction terms, to control for any residual imbalance after matching and to account for variation in baseline pageview levels. While this specification is consistent with best practices for doubly-robust estimation, it raises the question of whether the number of parameters is appropriate given the effective sample size at the species level. In the uplist magnitude model, for instance, 255 species support 29 fixed effects, yielding approximately nine species per parameter—near the recommended lower bound of linear mixed models with random slopes.

To assess whether our results are sensitive to covariate specification, we re-fitted all DiD models using only the variables that entered the matching procedure: the seven distance covariates (log-transformed body mass, absolute latitude, web capacity index, years known to science, and indicators for Use-Food, Use-Pet, and Use-Sport) plus population trend. The matching procedure itself was held constant (identical pairs, identical exact match constraints); only the outcome model specification was altered. This reduced the fixed effects from 27-30 to 17-19 per model, yielding a more conservative ratio of approximately 21 species per parameter in the tightest (uplist magnitude) specification. The meta-regression analysis (Section 3.5) was left unchanged, as it already operates at the species level with an appropriate parameter-to-observation ratio.

The simplified models produced DiD estimates that were virtually identical to the full specification. The uplist immediate effect remained significant (0.066, p = 0.025 vs. 0.067, p = 0.023 in the full model), and the downlist estimates were unchanged. The full DiD model with a three-way interaction with change authenticity remained consistent: the uplist immediate effect was significant (post_event × Treated_Uplist: 0.130, p = 0.006), and the change authenticity interaction terms were non-significant. Model convergence was clean in all specifications, with no singularity or rank deficiency warnings in any of the principal models.

This robustness demonstrates that our results are not driven by the inclusion of a large set of auxiliary covariates. Additionally, it confirms that the matching procedure alone achieves sufficient covariate balance that the outcome model specification exerts minimal influence on the estimated treatment effects. The simplified model can be viewed as a more parsimonious alternative; we report the full specification in the main text to maintain consistency with the doubly-robust DiD framework and to provide the fullest possible control for residual confounding.

**Supplementary Results**

1. **Consistency of drivers across major bird orders**

To test the generality of our findings, we repeated the baseline interest analyses on four major avian orders with distinct ecological and cultural profiles: Passeriformes (N = 6315, the largest avian order), Psittaciformes (parrots, N = 384), Accipitriformes (most raptors, N = 248), and Charadriiformes (shorebirds, gulls and auks, N = 366).

Despite many consistent factors across these diverse clades, the analyses revealed several clade-specific patterns. For Passeriformes (Figure S3), the drivers of interest closely mirrored the global patterns, though the effect of body mass was notably weaker for public (Estimate ± SE: 0.12 ± 0.01, p < 0.01) interest, and became non-significant for scientific interest (p = 0.82). Notably, use in other forms (e.g., medicines, handicrafts, etc.) emerged as a strong predictor of both public and scientific interest for Passeriformes (public: 0.52 ± 0.11, scientific: 0.56 ± 0.18, both p < 0.01).

For Accipitriformes (Figure S4), increasing populations (public: 0.93 ± 0.16, scientific: 0.81 ± 0.26, both p < 0.01) and web capacity index (public: 0.50 ± 0.06, scientific: 0.73 ± 0.09, both p < 0.01) were particularly strong predictors of both public and scientific interest. Those used as pets attracted significantly more scientific interest (0.67 ± 0.16, p < 0.01), where the effect size being substantially larger than in any other group.

The models for Psittaciformes highlighted the unique importance of charisma (Figure S5). Larger relative tail length was a significant predictor of public interest for this group (0.23 ± 0.07, p < 0.01), and generation length was a strong predictor for both domains (public: 0.33 ± 0.09, scientific: 0.33 ± 0.11, both p < 0.01). Notably, being threatened (public: 0.27 ± 0.12, p = 0.02; scientific: 0.66 ± 0.15, p < 0.01) and used as food (public: 0.61 ± 0.12, scientific: 0.73 ± 0.15, both p < 0.01) were the strongest factors of both public and scientific interest for Psittaciformes.

For Charadriiformes, factors related to familiarity were the most important drivers of public attention (Figure S6). Compared with other taxa, non-migrants in Charafriiformes received far less attention from both public and scientists (public: -0.30 ± 0.08, scientific: -0.54 ± 0.12, both p < 0.01). Besides, being threatened was not a significant driver of public interest for Charadriiformes (p = 0.10), and its effect size was also less pronounced for scientific interest (0.35 ± 0.15, p = 0.02).

**Figure S1. Comparative analysis of the drivers of public interest across major avian orders.**

The figure displays the standardized fixed-effect coefficients (Estimated Effect Size) and their 95% confidence intervals from five separate Generalized Linear Mixed Models (GLMMs) predicting public interest (number of Wikipedia pageviews). Each panel represents a single predictor variable, allowing for direct comparison of its effect size across the different taxonomic subsets: the global All Species model (N =10,509), and the subset models for Passeriformes (N = 6,315), Accipitriformes (N = 248), Charadriiformes (N = 366), and Psittaciformes (N = 384). Filled points indicate a statistically significant effect (p < 0.05), while hollow points indicate a non-significant effect. The vertical dashed line at zero represents a null effect. Full model outputs are provided in Table S2.

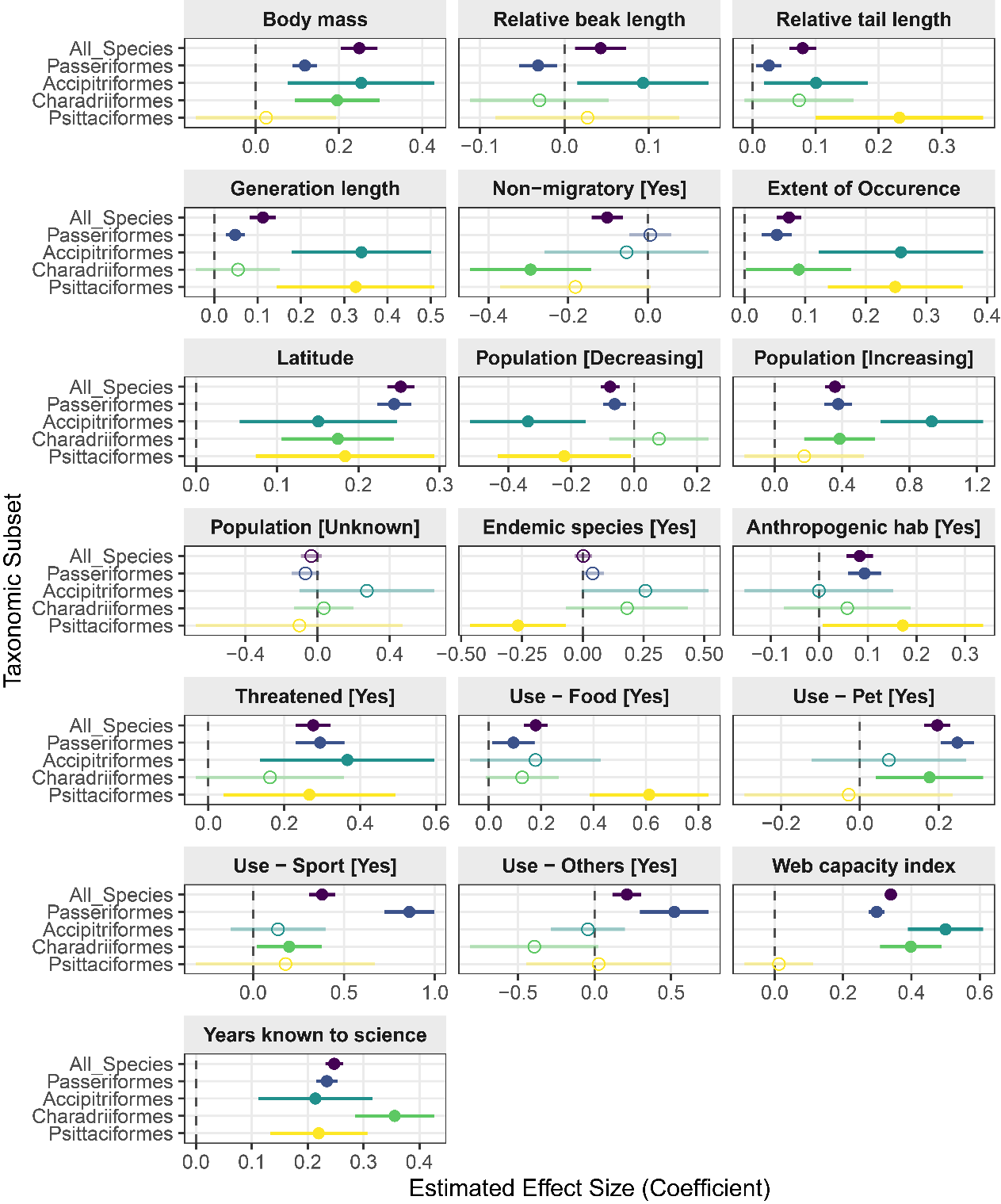

**Figure S2. Comparative analysis of the drivers of science interest across major avian orders**

The figure displays the standardized fixed-effect coefficients (Estimated Effect Size) and their 95% confidence intervals from five separate Generalized Linear Mixed Models (GLMMs) predicting scientific interest (number of Web of Science articles). Each panel represents a single predictor variable, allowing for direct comparison of its effect size across the different taxonomic subsets: the global All Species model (N =10,509), and the subset models for Passeriformes (N = 6,315), Accipitriformes (N = 248), Charadriiformes (N = 366), and Psittaciformes (N = 384).. Filled points indicate a statistically significant effect (p < 0.05), while hollow points indicate a non-significant effect. The vertical dashed line at zero represents a null effect. Full model outputs are provided in Table S2.

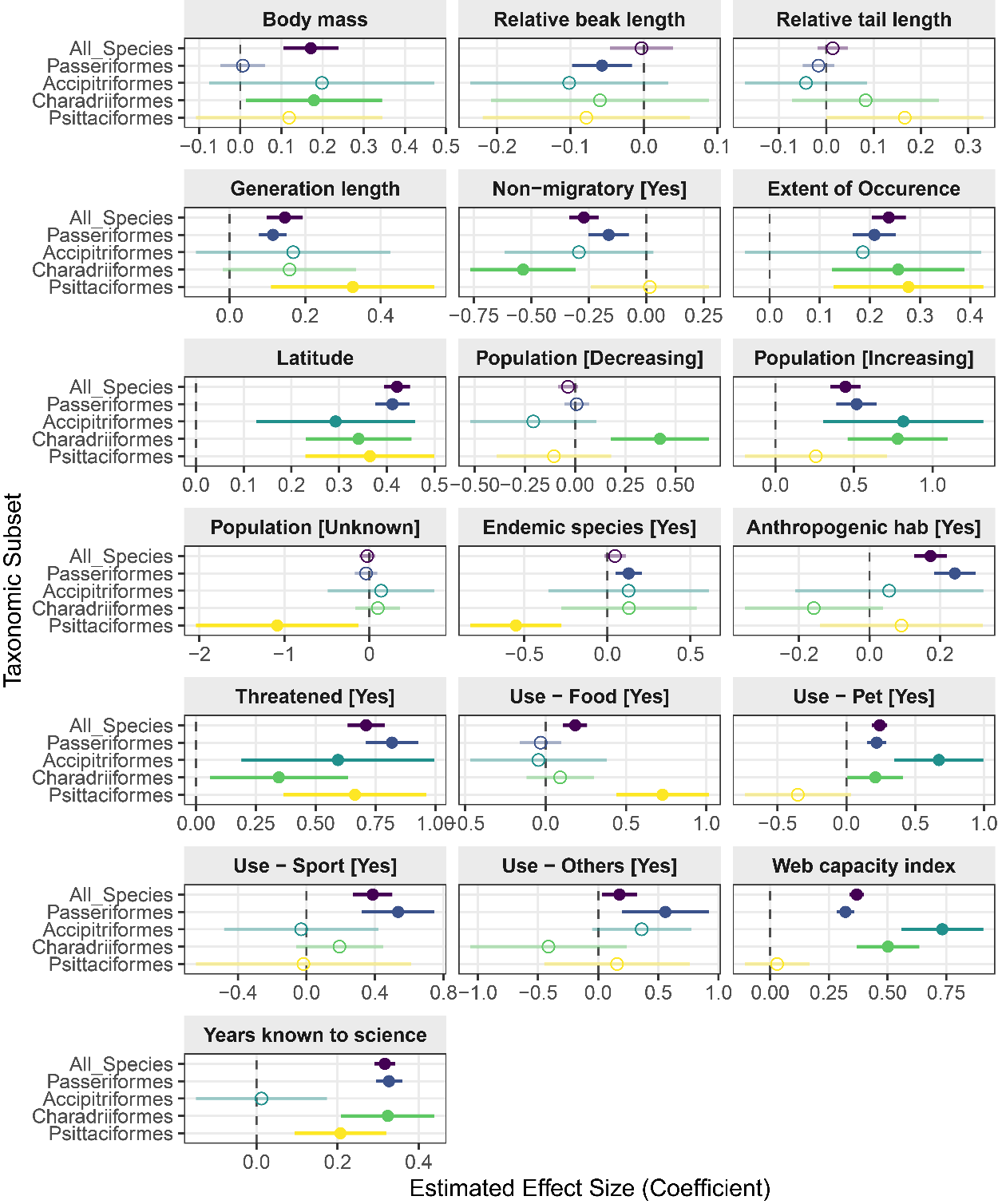

**Figure S3 Comparative profiles illustrating the drivers of public and scientific interest for Passeriformes.**

The figure displays the standardized fixed-effect coefficients from the baseline interest Generalized Linear Mixed Models (GLMMs), run independently for Passeriformes (N = 6315). The left radar plot shows the drivers for Public Interest (blue), and the right plot shows the drivers for Scientific Interest (orange), with the global model based on all bird species as reference (in grey). Each axis represents a predictor variable included in the models. The position of the point along each axis indicates the standardized coefficient (effect size) for that variable. The concentric dotted circles represent the scale of the effect size, with the circle at 0 indicating a null effect. Points located outside this zero-line represent positive effects (i.e., the trait is associated with higher interest), while points inside the circle represent negative effects. To highlight the most robust findings, predictors with a statistically significant effect (p < 0.05) are shown with a solid, high-opacity spoke and point, while non-significant predictors are faded to a lower opacity. ‘Stable’ was set as the reference level for Population trend. Full model outputs are provided in Table S2.

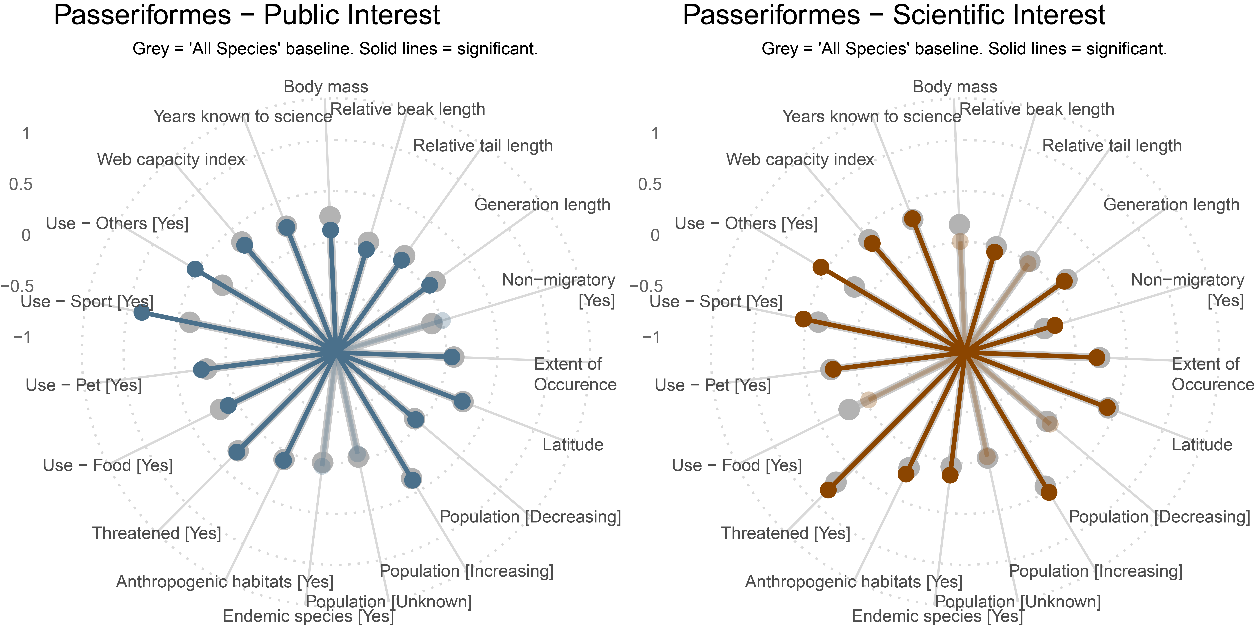

**Figure S4 Comparative profiles illustrating the drivers of public and scientific interest for Accipitriformes.**

The figure displays the standardized fixed-effect coefficients from the baseline interest Generalized Linear Mixed Models (GLMMs), run independently for Accipitriformes (N = 248). The left radar plot shows the drivers for Public Interest (blue), and the right plot shows the drivers for Scientific Interest (orange), with the global model based on all bird species as reference (in grey). Each axis represents a predictor variable included in the models. The position of the point along each axis indicates the standardized coefficient (effect size) for that variable. The concentric dotted circles represent the scale of the effect size, with the circle at 0 indicating a null effect. Points located outside this zero-line represent positive effects (i.e., the trait is associated with higher interest), while points inside the circle represent negative effects. To highlight the most robust findings, predictors with a statistically significant effect (p < 0.05) are shown with a solid, high-opacity spoke and point, while non-significant predictors are faded to a lower opacity. ‘Stable’ was set as the reference level for Population trend. Full model outputs are provided in Table S2.

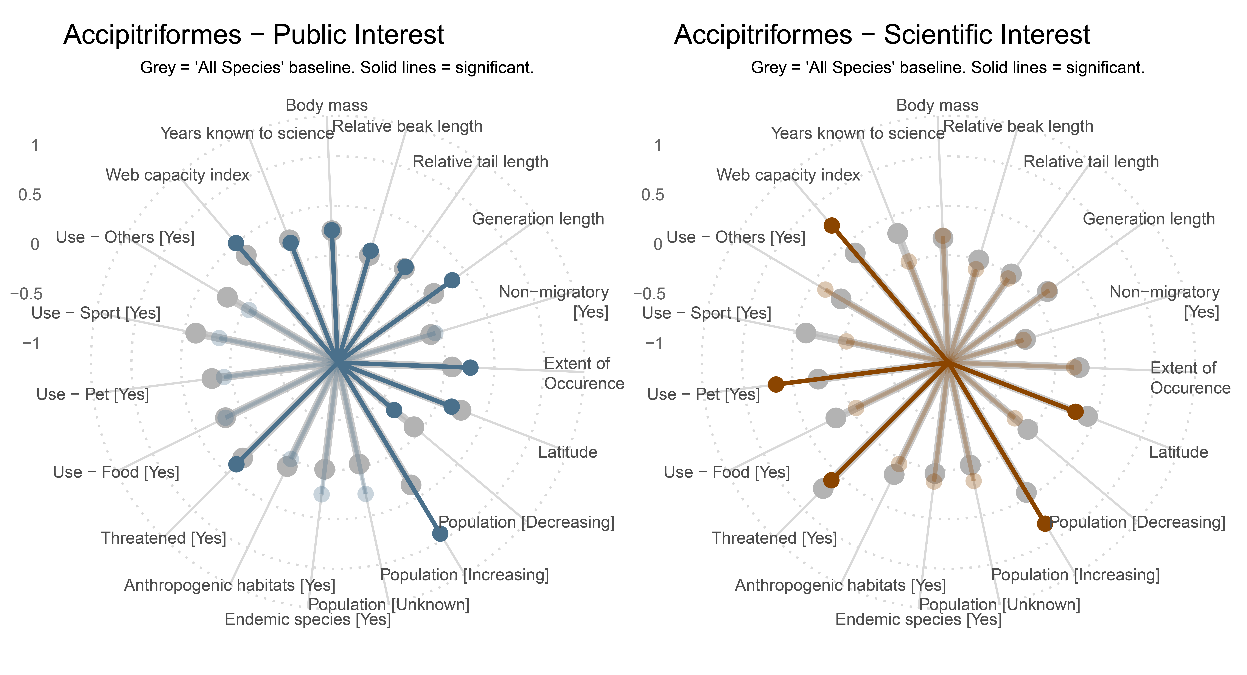

**Figure S5 Comparative profiles illustrating the drivers of public and scientific interest for Psittaciformes.**

The figure displays the standardized fixed-effect coefficients from the baseline interest Generalized Linear Mixed Models (GLMMs), run independently for Psittaciformes (N = 384). The left radar plot shows the drivers for Public Interest (blue), and the right plot shows the drivers for Scientific Interest (orange), with the global model based on all bird species as reference (in grey). Each axis represents a predictor variable included in the models. The position of the point along each axis indicates the standardized coefficient (effect size) for that variable. The concentric dotted circles represent the scale of the effect size, with the circle at 0 indicating a null effect. Points located outside this zero-line represent positive effects (i.e., the trait is associated with higher interest), while points inside the circle represent negative effects. To highlight the most robust findings, predictors with a statistically significant effect (p < 0.05) are shown with a solid, high-opacity spoke and point, while non-significant predictors are faded to a lower opacity. ‘Stable’ was set as the reference level for Population trend. Full model outputs are provided in Table S2.

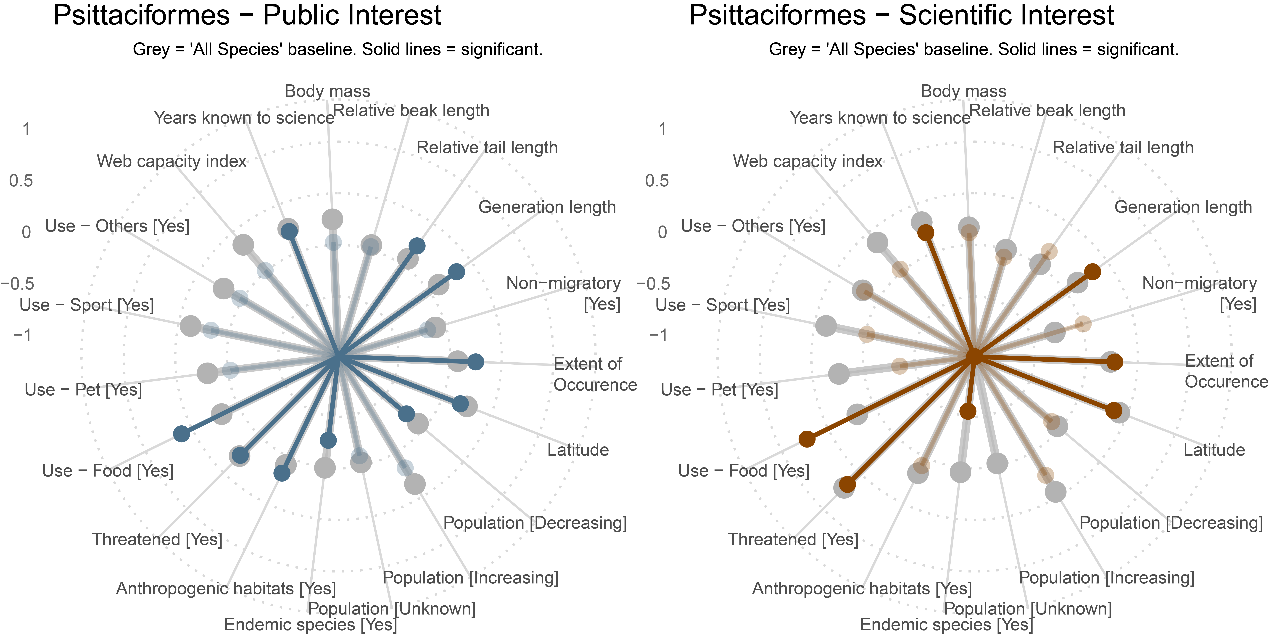

**Figure S6 Comparative profiles illustrating the drivers of public and scientific interest for Charadriiformes.**

The figure displays the standardized fixed-effect coefficients from the baseline interest Generalized Linear Mixed Models (GLMMs), run independently for Charadriiformes (N = 366). The left radar plot shows the drivers for Public Interest (blue), and the right plot shows the drivers for Scientific Interest (orange), with the global model based on all bird species as reference (in grey). Each axis represents a predictor variable included in the models. The position of the point along each axis indicates the standardized coefficient (effect size) for that variable. The concentric dotted circles represent the scale of the effect size, with the circle at 0 indicating a null effect. Points located outside this zero-line represent positive effects (i.e., the trait is associated with higher interest), while points inside the circle represent negative effects. To highlight the most robust findings, predictors with a statistically significant effect (p < 0.05) are shown with a solid, high-opacity spoke and point, while non-significant predictors are faded to a lower opacity. ‘Stable’ was set as the reference level for Population trend. Full model outputs are provided in Table S2.

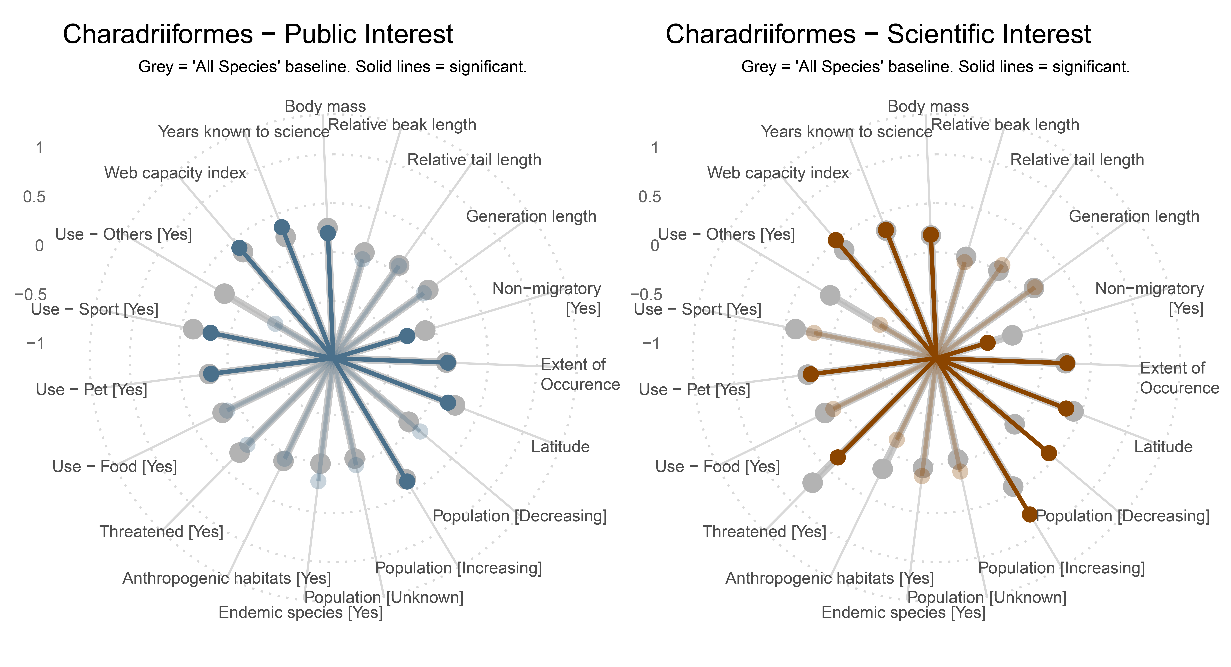

**Figure S7. Drivers of relative scientific and public interest.**

Forest plots summarize the estimated parameters (effect size mean ± 95% CI) based on Gaussian linear mixed models testing the relationship between residuals from the linear regression line in Figure 2 and the final set of predictors (marginal R^2^ = 0.094, N = 10509). Positive residuals indicate species with a greater popular than scientific interest, and negative values indicate the opposite way. Factor baselines: Population trend (Stable). Filled points indicate statistically significant effects (p < 0.05). Estimated regression parameters and p-values are in Table S3.

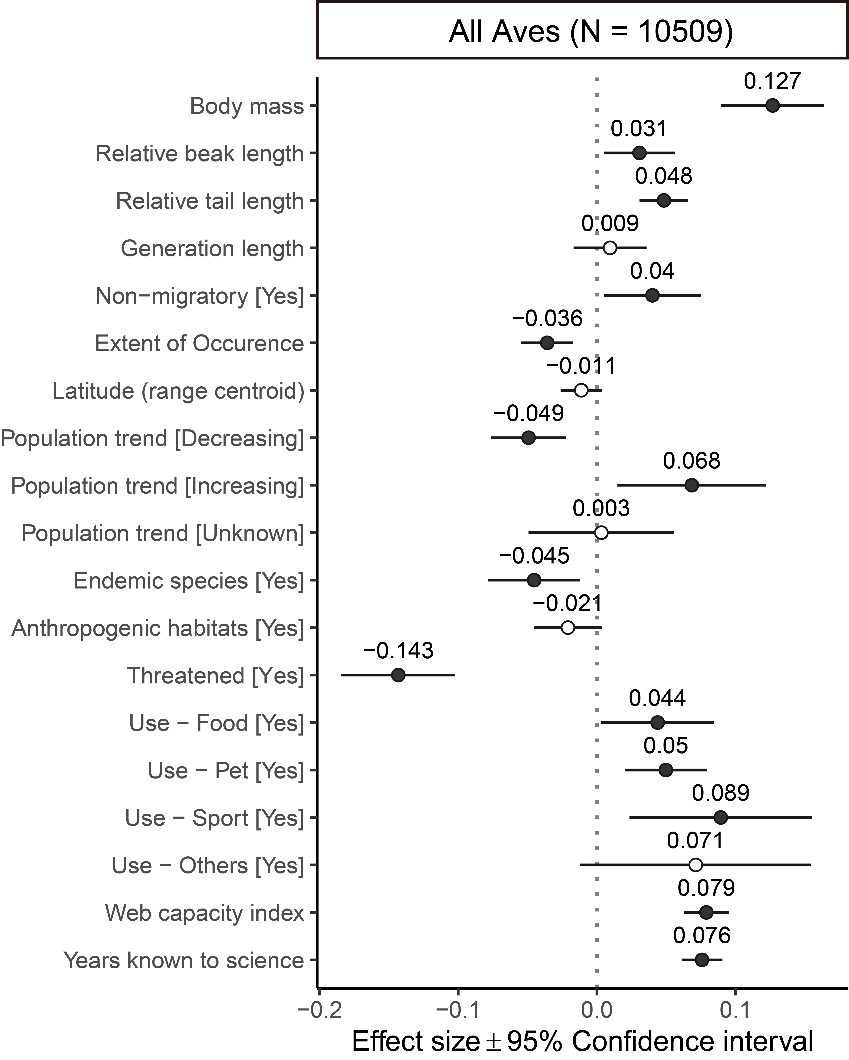

**Table S1.** Estimated regression parameters of the baseline public and scientific interest model for all bird species.

|  | Public interest model | | | |  | Scientific interest model | | | |
| --- | --- | --- | --- | --- | --- | --- | --- | --- | --- |
| Variable | Estimate | SE | z | p |  | Estimate | SE | z | p |
| Non-migratory [Yes] | -0.102 | 0.020 | -5.149 | <0.001 |  | -0.271 | 0.032 | -8.398 | <0.001 |
| Body mass | 0.249 | 0.022 | 11.092 | <0.001 |  | 0.172 | 0.034 | 5.019 | <0.001 |
| Extent of Occurrence | 0.073 | 0.010 | 7.050 | <0.001 |  | 0.238 | 0.017 | 13.731 | <0.001 |
| Latitude (range centroid) | 0.252 | 0.008 | 29.710 | <0.001 |  | 0.421 | 0.014 | 30.377 | <0.001 |
| Relative beak length | 0.043 | 0.016 | 2.771 | 0.006 |  | -0.003 | 0.022 | -0.151 | 0.880 |
| Relative tail length | 0.080 | 0.011 | 7.298 | <0.001 |  | 0.014 | 0.017 | 0.822 | 0.411 |
| Threatened [Yes] | 0.276 | 0.024 | 11.735 | <0.001 |  | 0.711 | 0.039 | 18.120 | <0.001 |
| Use - Food [Yes] | 0.179 | 0.023 | 7.637 | <0.001 |  | 0.184 | 0.038 | 4.843 | <0.001 |
| Use - Pet [Yes] | 0.196 | 0.017 | 11.529 | <0.001 |  | 0.242 | 0.028 | 8.640 | <0.001 |
| Use - Sport [Yes] | 0.380 | 0.037 | 10.285 | <0.001 |  | 0.387 | 0.058 | 6.624 | <0.001 |
| Use - Others [Yes] | 0.211 | 0.048 | 4.356 | <0.001 |  | 0.177 | 0.075 | 2.358 | 0.018 |
| Population trend [Unknown] | -0.036 | 0.030 | -1.205 | 0.228 |  | -0.026 | 0.049 | -0.519 | 0.604 |
| Population trend [Decreasing] | -0.076 | 0.015 | -5.025 | <0.001 |  | -0.036 | 0.026 | -1.391 | 0.164 |
| Population trend [Increasing] | 0.358 | 0.031 | 11.650 | <0.001 |  | 0.446 | 0.048 | 9.196 | <0.001 |
| Generation length | 0.112 | 0.015 | 7.266 | <0.001 |  | 0.146 | 0.024 | 6.017 | <0.001 |
| Endemic species [Yes] | 0.002 | 0.019 | 0.107 | 0.915 |  | 0.046 | 0.033 | 1.376 | 0.169 |

**Table S2.** Estimated regression parameters of the baseline public and scientific interest model for subset bird orders.

|  | Public interest model | | | |  | Scientific interest model | | | |
| --- | --- | --- | --- | --- | --- | --- | --- | --- | --- |
| Variable | Estimate | SE | z | p |  | Estimate | SE | z | p |
| **Passeriformes** |  |  |  |  |  |  |  |  |  |
| Non-migratory [Yes] | 0.006 | 0.027 | 0.218 | 0.827 |  | -0.163 | 0.045 | -3.664 | <0.001 |
| Body mass | 0.118 | 0.015 | 7.900 | <0.001 |  | 0.006 | 0.028 | 0.225 | 0.822 |
| Extent of Occurrence | 0.053 | 0.013 | 4.191 | <0.001 |  | 0.209 | 0.022 | 9.561 | <0.001 |
| Latitude (range centroid) | 0.244 | 0.011 | 22.684 | <0.001 |  | 0.412 | 0.018 | 22.633 | <0.001 |
| Relative beak length | -0.031 | 0.012 | -2.691 | 0.007 |  | -0.057 | 0.021 | -2.746 | 0.006 |
| Relative tail length | 0.026 | 0.010 | 2.562 | 0.010 |  | -0.017 | 0.017 | -0.986 | 0.324 |
| Threatened [Yes] | 0.294 | 0.033 | 8.998 | <0.001 |  | 0.819 | 0.056 | 14.751 | <0.001 |
| Use - Food [Yes] | 0.094 | 0.041 | 2.264 | 0.024 |  | -0.031 | 0.066 | -0.471 | 0.637 |
| Use - Pet [Yes] | 0.247 | 0.021 | 11.550 | <0.001 |  | 0.219 | 0.036 | 6.060 | <0.001 |
| Use - Sport [Yes] | 0.860 | 0.071 | 12.170 | <0.001 |  | 0.535 | 0.109 | 4.929 | <0.001 |
| Use - Others [Yes] | 0.521 | 0.115 | 4.547 | <0.001 |  | 0.558 | 0.185 | 3.021 | 0.003 |
| Population trend [Unknown] | -0.068 | 0.039 | -1.722 | 0.085 |  | -0.039 | 0.067 | -0.580 | 0.562 |
| Population trend [Decreasing] | -0.062 | 0.018 | -3.372 | 0.001 |  | 0.007 | 0.031 | 0.235 | 0.814 |
| Population trend [Increasing] | 0.377 | 0.042 | 9.060 | <0.001 |  | 0.517 | 0.066 | 7.844 | <0.001 |
| Generation length | 0.048 | 0.011 | 4.321 | <0.001 |  | 0.114 | 0.019 | 6.077 | <0.001 |
| Endemic species [Yes] | 0.041 | 0.023 | 1.768 | 0.077 |  | 0.128 | 0.041 | 3.121 | 0.002 |
| Anthropogenic habitats [Yes] | 0.093 | 0.017 | 5.398 | <0.001 |  | 0.242 | 0.030 | 8.029 | <0.001 |
| Web capacity index | 0.298 | 0.012 | 24.896 | <0.001 |  | 0.321 | 0.019 | 16.579 | <0.001 |
| Years known to science | 0.234 | 0.010 | 23.867 | <0.001 |  | 0.327 | 0.017 | 19.616 | <0.001 |
| **Accipitriformes** |  |  |  |  |  |  |  |  |  |
| Non-migratory [Yes] | -0.054 | 0.105 | -0.510 | 0.610 |  | -0.293 | 0.166 | -1.771 | 0.077 |
| Body mass | 0.253 | 0.090 | 2.819 | 0.005 |  | 0.198 | 0.140 | 1.422 | 0.155 |
| Extent of Occurence | 0.258 | 0.069 | 3.726 | <0.001 |  | 0.186 | 0.120 | 1.546 | 0.122 |
| Latitude (range centroid) | 0.151 | 0.049 | 3.053 | 0.002 |  | 0.293 | 0.085 | 3.449 | 0.001 |
| Relative beak length | 0.093 | 0.040 | 2.345 | 0.019 |  | -0.102 | 0.069 | -1.482 | 0.138 |
| Relative tail length | 0.101 | 0.042 | 2.400 | 0.016 |  | -0.043 | 0.066 | -0.652 | 0.514 |
| Threatened [Yes] | 0.366 | 0.117 | 3.126 | 0.002 |  | 0.593 | 0.206 | 2.883 | 0.004 |
| Use - Food [Yes] | 0.178 | 0.127 | 1.401 | 0.161 |  | -0.045 | 0.216 | -0.207 | 0.836 |
| Use - Pet [Yes] | 0.073 | 0.100 | 0.734 | 0.463 |  | 0.672 | 0.165 | 4.078 | <0.001 |
| Use - Sport [Yes] | 0.137 | 0.133 | 1.029 | 0.304 |  | -0.031 | 0.230 | -0.136 | 0.892 |
| Use - Others [Yes] | -0.044 | 0.123 | -0.353 | 0.724 |  | 0.361 | 0.210 | 1.720 | 0.085 |
| Population trend [Unknown] | 0.273 | 0.191 | 1.427 | 0.154 |  | 0.139 | 0.321 | 0.432 | 0.665 |
| Population trend [Decreasing] | -0.337 | 0.094 | -3.603 | <0.001 |  | -0.208 | 0.160 | -1.302 | 0.193 |
| Population trend [Increasing] | 0.931 | 0.156 | 5.978 | <0.001 |  | 0.814 | 0.260 | 3.135 | 0.002 |
| Generation length | 0.340 | 0.082 | 4.139 | <0.001 |  | 0.168 | 0.131 | 1.280 | 0.200 |
| Endemic species [Yes] | 0.257 | 0.133 | 1.933 | 0.053 |  | 0.127 | 0.247 | 0.516 | 0.606 |
| Anthropogenic habitats [Yes] | -0.001 | 0.078 | -0.007 | 0.995 |  | 0.056 | 0.136 | 0.409 | 0.683 |
| Web capacity index | 0.500 | 0.056 | 8.896 | <0.001 |  | 0.734 | 0.089 | 8.239 | <0.001 |
| Years known to science | 0.214 | 0.052 | 4.094 | <0.001 |  | 0.012 | 0.082 | 0.150 | 0.880 |
| **Psittaciformes** |  |  |  |  |  |  |  |  |  |
| Non-migratory [Yes] | -0.182 | 0.096 | -1.891 | 0.059 |  | 0.016 | 0.131 | 0.120 | 0.904 |
| Body mass | 0.025 | 0.086 | 0.288 | 0.773 |  | 0.119 | 0.115 | 1.028 | 0.304 |
| Extent of Occurrence | 0.248 | 0.057 | 4.382 | <0.001 |  | 0.277 | 0.076 | 3.635 | <0.001 |
| Latitude (range centroid) | 0.184 | 0.056 | 3.269 | 0.001 |  | 0.365 | 0.069 | 5.281 | <0.001 |
| Relative beak length | 0.027 | 0.056 | 0.492 | 0.623 |  | -0.078 | 0.072 | -1.088 | 0.276 |
| Relative tail length | 0.233 | 0.068 | 3.440 | 0.001 |  | 0.166 | 0.085 | 1.958 | 0.050 |
| Threatened [Yes] | 0.266 | 0.116 | 2.304 | 0.021 |  | 0.664 | 0.153 | 4.352 | <0.001 |
| Use - Food [Yes] | 0.612 | 0.116 | 5.269 | <0.001 |  | 0.727 | 0.147 | 4.937 | <0.001 |
| Use - Pet [Yes] | -0.028 | 0.135 | -0.210 | 0.834 |  | -0.353 | 0.197 | -1.790 | 0.073 |
| Use - Sport [Yes] | 0.179 | 0.251 | 0.710 | 0.478 |  | -0.018 | 0.320 | -0.055 | 0.956 |
| Use - Others [Yes] | 0.027 | 0.241 | 0.112 | 0.911 |  | 0.157 | 0.309 | 0.507 | 0.612 |
| Population trend [Unknown] | -0.101 | 0.293 | -0.346 | 0.730 |  | -1.084 | 0.488 | -2.222 | 0.026 |
| Population trend [Decreasing] | -0.222 | 0.108 | -2.050 | 0.040 |  | -0.106 | 0.146 | -0.726 | 0.468 |
| Population trend [Increasing] | 0.175 | 0.181 | 0.968 | 0.333 |  | 0.257 | 0.231 | 1.115 | 0.265 |
| Generation length | 0.327 | 0.093 | 3.515 | <0.001 |  | 0.326 | 0.110 | 2.951 | 0.003 |
| Endemic species [Yes] | -0.267 | 0.100 | -2.662 | 0.008 |  | -0.550 | 0.140 | -3.929 | <0.001 |
| Anthropogenic habitats [Yes] | 0.172 | 0.084 | 2.039 | 0.041 |  | 0.091 | 0.118 | 0.770 | 0.442 |
| Web capacity index | 0.012 | 0.051 | 0.243 | 0.808 |  | 0.030 | 0.070 | 0.422 | 0.673 |
| Years known to science | 0.220 | 0.044 | 4.942 | <0.001 |  | 0.207 | 0.057 | 3.608 | <0.001 |
| **Charadriiformes** |  |  |  |  |  |  |  |  |  |
| Non-migratory [Yes] | -0.294 | 0.077 | -3.814 | <0.001 |  | -0.536 | 0.117 | -4.570 | <0.001 |
| Body mass | 0.195 | 0.052 | 3.756 | <0.001 |  | 0.179 | 0.084 | 2.124 | 0.034 |
| Extent of Occurrence | 0.089 | 0.044 | 2.018 | 0.044 |  | 0.256 | 0.067 | 3.799 | <0.001 |
| Latitude (range centroid) | 0.175 | 0.035 | 4.952 | <0.001 |  | 0.341 | 0.057 | 6.015 | <0.001 |
| Relative beak length | -0.030 | 0.042 | -0.706 | 0.480 |  | -0.060 | 0.076 | -0.791 | 0.429 |
| Relative tail length | 0.074 | 0.044 | 1.680 | 0.093 |  | 0.083 | 0.079 | 1.047 | 0.295 |
| Threatened [Yes] | 0.163 | 0.100 | 1.631 | 0.103 |  | 0.346 | 0.147 | 2.355 | 0.019 |
| Use - Food [Yes] | 0.127 | 0.071 | 1.784 | 0.074 |  | 0.090 | 0.106 | 0.848 | 0.397 |
| Use - Pet [Yes] | 0.176 | 0.069 | 2.549 | 0.011 |  | 0.210 | 0.103 | 2.035 | 0.042 |
| Use - Sport [Yes] | 0.198 | 0.091 | 2.178 | 0.029 |  | 0.193 | 0.129 | 1.487 | 0.137 |
| Use - Others [Yes] | -0.393 | 0.213 | -1.840 | 0.066 |  | -0.413 | 0.331 | -1.247 | 0.212 |
| Population trend [Unknown] | 0.034 | 0.084 | 0.404 | 0.686 |  | 0.100 | 0.134 | 0.743 | 0.457 |
| Population trend [Decreasing] | 0.079 | 0.081 | 0.978 | 0.328 |  | 0.423 | 0.124 | 3.399 | 0.001 |
| Population trend [Increasing] | 0.385 | 0.108 | 3.578 | <0.001 |  | 0.778 | 0.163 | 4.782 | <0.001 |
| Generation length | 0.055 | 0.050 | 1.105 | 0.269 |  | 0.158 | 0.090 | 1.757 | 0.079 |
| Endemic species [Yes] | 0.182 | 0.128 | 1.421 | 0.155 |  | 0.130 | 0.208 | 0.624 | 0.533 |
| Anthropogenic habitats [Yes] | 0.058 | 0.066 | 0.871 | 0.384 |  | -0.158 | 0.100 | -1.580 | 0.114 |
| Web capacity index | 0.398 | 0.046 | 8.687 | <0.001 |  | 0.502 | 0.068 | 7.361 | <0.001 |
| Years known to science | 0.356 | 0.036 | 9.774 | <0.001 |  | 0.324 | 0.059 | 5.507 | <0.001 |

**Table S3.** Estimated regression parameters of the relative scientific and public interest model across all bird species.

| Variable | Estimate | SE | z | p |
| --- | --- | --- | --- | --- |
| Non-migratory [Yes] | 0.040 | 0.018 | 2.262 | 0.024 |
| Body mass | 0.127 | 0.019 | 6.692 | <0.001 |
| Extent of Occurence | -0.036 | 0.010 | -3.760 | <0.001 |
| Latitude (range centroid) | -0.011 | 0.008 | -1.476 | 0.140 |
| Relative beak length | 0.031 | 0.013 | 2.350 | 0.019 |
| Relative tail length | 0.048 | 0.009 | 5.377 | <0.001 |
| Threatened [Yes] | -0.143 | 0.021 | -6.826 | <0.001 |
| Use - Food [Yes] | 0.044 | 0.021 | 2.099 | 0.036 |
| Use - Pet [Yes] | 0.050 | 0.015 | 3.311 | 0.001 |
| Use - Sport [Yes] | 0.089 | 0.034 | 2.659 | 0.008 |
| Use - Others [Yes] | 0.071 | 0.042 | 1.679 | 0.093 |
| Population trend [Unknown] | 0.003 | 0.027 | 0.120 | 0.905 |
| Population trend [Decreasing] | -0.049 | 0.014 | -3.571 | <0.001 |
| Population trend [Increasing] | 0.068 | 0.027 | 2.498 | 0.012 |
| Generation length | 0.009 | 0.013 | 0.706 | 0.480 |
| Endemic species [Yes] | -0.045 | 0.017 | -2.680 | 0.007 |
| Anthropogenic habitats [Yes] | -0.021 | 0.012 | -1.677 | 0.094 |
| Web capacity index | 0.079 | 0.008 | 9.647 | <0.001 |
| Years known to science | 0.076 | 0.007 | 10.112 | <0.001 |

**Table S4.** Full fixed-effects results from the Difference-in-Differences (DiD) Linear Mixed-Effects Models. The key DiD estimates are the interaction terms representing the immediate (Post-Event x Group) and trend (Time Since Event x Group) effects.

| Model | Term | Estimate | Std Error | t | p value |
| --- | --- | --- | --- | --- | --- |
| Model A: Primary DiD (all changes) (N = 6624 obs, 736 species) | | | | | |
|  | Time (baseline trend) | 0.074 | 0.006 | 12.46 | <0.001*** |
|  | Body mass | 0.217 | 0.040 | 5.47 | <0.001*** |
|  | Absolute latitude | 0.210 | 0.028 | 7.57 | <0.001*** |
|  | Web Capacity Index | 0.129 | 0.031 | 4.18 | <0.001*** |
|  | Years known to science | 0.195 | 0.031 | 6.31 | <0.001*** |
|  | Use-Food | 0.129 | 0.076 | 1.69 | 0.091. |
|  | Use-Pet | 0.475 | 0.060 | 7.91 | <0.001*** |
|  | Use-Sport | 0.535 | 0.145 | 3.68 | <0.001*** |
|  | Population trend-Decreasing | -0.212 | 0.071 | -3.01 | 0.003** |
|  | Population trend-Increasing | 0.127 | 0.131 | 0.97 | 0.332 |
|  | Extent of Occurrence | 0.058 | 0.033 | 1.73 | 0.084. |
|  | Generation length | 0.114 | 0.036 | 3.15 | 0.002** |
|  | Relative beak length | 0.002 | 0.025 | 0.07 | 0.947 |
|  | Relative tail length | 0.016 | 0.025 | 0.64 | 0.524 |
|  | Use-Other | 0.424 | 0.179 | 2.37 | 0.018* |
|  | Endemic species | 0.032 | 0.059 | 0.54 | 0.588 |
|  | Anthropogenic habitat use | -0.021 | 0.052 | -0.41 | 0.681 |
|  | Non-migrant status [Yes] | -0.174 | 0.078 | -2.22 | 0.027* |
|  | Post-event (control group) | 0.076 | 0.016 | 4.79 | <0.001*** |
|  | Group-Treated Downlist | 0.016 | 0.070 | 0.23 | 0.818 |
|  | Group-Treated Uplist | -0.024 | 0.089 | -0.27 | 0.787 |
|  | Time since event (control group) | 0.016 | 0.007 | 2.41 | 0.016* |
|  | Post-event × Downlist | -0.016 | 0.024 | -0.69 | 0.491 |
|  | Post-event × Uplist | 0.067 | 0.030 | 2.27 | 0.023* |
|  | Time-since-event × Downlist | 0.011 | 0.010 | 1.16 | 0.244 |
|  | Time-since-event × Uplist | -0.002 | 0.012 | -0.18 | 0.858 |
| Model B: Uplist by magnitude (N = 2295 obs, 255 species) | | | | | |
|  | Time (baseline trend) | 0.101 | 0.014 | 7.23 | <0.001*** |
|  | Body mass | 0.194 | 0.076 | 2.57 | 0.011* |
|  | Absolute latitude | 0.200 | 0.049 | 4.04 | <0.001*** |
|  | Web Capacity Index | 0.249 | 0.052 | 4.74 | <0.001*** |
|  | Years known to science | 0.265 | 0.059 | 4.49 | <0.001*** |
|  | Extent of Occurrence | 0.053 | 0.066 | 0.80 | 0.424 |
|  | Generation length | 0.191 | 0.065 | 2.92 | 0.004** |
|  | Relative beak length | -0.033 | 0.045 | -0.73 | 0.467 |
|  | Relative tail length | 0.035 | 0.045 | 0.79 | 0.431 |
|  | Use-Food | 0.051 | 0.124 | 0.42 | 0.677 |
|  | Use-Pet | 0.487 | 0.100 | 4.89 | <0.001*** |
|  | Use-Sport | 0.505 | 0.205 | 2.46 | 0.015* |
|  | Use-Other | 0.411 | 0.257 | 1.60 | 0.111 |
|  | Endemic species | -0.056 | 0.111 | -0.50 | 0.616 |
|  | Anthropogenic habitat use | -0.064 | 0.091 | -0.71 | 0.478 |
|  | Non-migrant status-Yes | -0.076 | 0.135 | -0.56 | 0.573 |
|  | Population trend-Decreasing | -0.157 | 0.478 | -0.33 | 0.743 |
|  | Population trend-Increasing | 0.235 | 0.683 | 0.34 | 0.731 |
|  | Post-event (control group) | 0.050 | 0.032 | 1.56 | 0.119 |
|  | Magnitude-High | 0.086 | 0.186 | 0.46 | 0.644 |
|  | Magnitude-Low | -0.011 | 0.167 | -0.07 | 0.948 |
|  | Magnitude-Moderate | 0.121 | 0.165 | 0.73 | 0.463 |
|  | Time since event (control group) | 0.010 | 0.013 | 0.76 | 0.444 |
|  | Post-event × High magnitude | 0.095 | 0.063 | 1.51 | 0.132 |
|  | Post-event × Low magnitude | 0.067 | 0.062 | 1.09 | 0.276 |
|  | Post-event × Moderate magnitude | 0.033 | 0.061 | 0.55 | 0.585 |
|  | Time-since-event × High magnitude | 0.003 | 0.025 | 0.14 | 0.893 |
|  | Time-since-event × Low magnitude | -0.013 | 0.025 | -0.53 | 0.597 |
|  | Time-since-event × Moderate magnitude | -0.019 | 0.025 | -0.77 | 0.441 |
| Model D: Downlist by magnitude (N = 3384 obs, 376 species) | | | | | |
|  | Time (baseline trend) | 0.070 | 0.007 | 10.28 | <0.001*** |
|  | Body mass | 0.156 | 0.050 | 3.09 | 0.002** |
|  | Absolute latitude | 0.241 | 0.038 | 6.34 | <0.001*** |
|  | Web Capacity Index | 0.013 | 0.041 | 0.31 | 0.758 |
|  | Years known to science | 0.100 | 0.039 | 2.56 | 0.011* |
|  | Extent of Occurrence | 0.053 | 0.046 | 1.17 | 0.243 |
|  | Generation length | 0.117 | 0.049 | 2.38 | 0.018* |
|  | Relative beak length | 0.087 | 0.035 | 2.50 | 0.013* |
|  | Relative tail length | 0.003 | 0.033 | 0.08 | 0.939 |
|  | Use-Food | 0.287 | 0.108 | 2.65 | 0.008** |
|  | Use-Pet | 0.447 | 0.080 | 5.57 | <0.001*** |
|  | Use-Sport | 0.652 | 0.238 | 2.74 | 0.006** |
|  | Use-Other | 0.488 | 0.211 | 2.32 | 0.021* |
|  | Endemic species | -0.008 | 0.078 | -0.10 | 0.921 |
|  | Anthropogenic habitat use | 0.032 | 0.070 | 0.45 | 0.650 |
|  | Non-migrant status-Yes | -0.073 | 0.122 | -0.60 | 0.546 |
|  | Population trend-Decreasing | -0.051 | 0.125 | -0.41 | 0.684 |
|  | Population trend-Increasing | 0.081 | 0.264 | 0.31 | 0.760 |
|  | Post-event (control group) | 0.078 | 0.021 | 3.76 | <0.001*** |
|  | Magnitude-High | -0.052 | 0.141 | -0.37 | 0.714 |
|  | Magnitude-Low | -0.172 | 0.107 | -1.60 | 0.109 |
|  | Magnitude-Moderate | 0.015 | 0.112 | 0.13 | 0.895 |
|  | Time since event (control group) | 0.001 | 0.009 | 0.07 | 0.945 |
|  | Post-event × High magnitude | -0.016 | 0.046 | -0.34 | 0.731 |
|  | Post-event × Low magnitude | 0.017 | 0.036 | 0.48 | 0.633 |
|  | Post-event × Moderate magnitude | -0.043 | 0.038 | -1.13 | 0.257 |
|  | Time-since-event × High magnitude | 0.030 | 0.018 | 1.66 | 0.097. |
|  | Time-since-event × Low magnitude | 0.019 | 0.015 | 1.27 | 0.205 |
|  | Time-since-event × Moderate magnitude | 0.021 | 0.016 | 1.35 | 0.176 |
| Model C: Genuine changes only (N = 6345 obs, 705 species) | | | | | |
|  | Time (baseline trend) | 0.074 | 0.006 | 11.97 | <0.001*** |
|  | Body mass | 0.307 | 0.031 | 9.75 | <0.001*** |
|  | Absolute latitude | 0.218 | 0.027 | 8.12 | <0.001*** |
|  | Web Capacity Index | 0.170 | 0.029 | 5.90 | <0.001*** |
|  | Years known to science | 0.202 | 0.029 | 6.96 | <0.001*** |
|  | Use-Food | 0.149 | 0.077 | 1.93 | 0.054. |
|  | Use-Pet | 0.505 | 0.061 | 8.31 | <0.001*** |
|  | Use-Sport | 0.456 | 0.146 | 3.11 | 0.002** |
|  | Population trend-Decreasing | -0.218 | 0.071 | -3.08 | 0.002** |
|  | Population trend-Increasing | 0.194 | 0.133 | 1.46 | 0.146 |
|  | Post-event (control group) | 0.080 | 0.016 | 4.91 | <0.001*** |
|  | Group-Treated Downlist | -0.005 | 0.072 | -0.06 | 0.948 |
|  | Group-Treated Uplist | -0.003 | 0.090 | -0.03 | 0.973 |
|  | Time since event (control group) | 0.015 | 0.007 | 2.19 | 0.029* |
|  | Post-event × Downlist | -0.021 | 0.025 | -0.84 | 0.401 |
|  | Post-event × Uplist | 0.064 | 0.030 | 2.11 | 0.034* |
|  | Time-since-event × Downlist | 0.015 | 0.010 | 1.48 | 0.139 |
|  | Time-since-event × Uplist | -0.002 | 0.012 | -0.13 | 0.893 |
| Model E: Non-genuine changes only (N = 6597 obs, 733 species) | | | | | |
|  | Time (baseline trend) | 0.074 | 0.006 | 12.37 | <0.001*** |
|  | Body mass | 0.323 | 0.032 | 10.24 | <0.001*** |
|  | Absolute latitude | 0.241 | 0.027 | 8.98 | <0.001*** |
|  | Web Capacity Index | 0.162 | 0.029 | 5.66 | <0.001*** |
|  | Years known to science | 0.211 | 0.029 | 7.28 | <0.001*** |
|  | Use-Food | 0.119 | 0.077 | 1.55 | 0.123 |
|  | Use-Pet | 0.517 | 0.060 | 8.59 | <0.001*** |
|  | Use-Sport | 0.439 | 0.146 | 3.01 | 0.003** |
|  | Population trend-Decreasing | -0.190 | 0.070 | -2.73 | 0.006** |
|  | Population trend-Increasing | 0.177 | 0.136 | 1.30 | 0.194 |
|  | Post-event (control group) | 0.077 | 0.016 | 4.88 | <0.001*** |
|  | Group-Treated Downlist | 0.008 | 0.070 | 0.12 | 0.905 |
|  | Group-Treated Uplist | -0.016 | 0.090 | -0.18 | 0.860 |
|  | Time since event (control group) | 0.017 | 0.007 | 2.53 | 0.011* |
|  | Post-event × Downlist | -0.017 | 0.024 | -0.70 | 0.483 |
|  | Post-event × Uplist | 0.066 | 0.030 | 2.22 | 0.026* |
|  | Time-since-event × Downlist | 0.011 | 0.010 | 1.15 | 0.251 |
|  | Time-since-event × Uplist | -0.003 | 0.012 | -0.27 | 0.789 |

**Table S5.** Full results from the Weighted Least Squares (WLS) meta-regression models. Models were fitted for the Overall sample (and a sensitivity check), the uplist and downlist subset, and were weighted by the inverse variance of the effect size estimates. Both uplist and downlist model are were not statistically significant at the α = 0.05 level (Uplist: F(21,108) = 1.49, p = 0.096, Adj. R² = 0.074; Downlist: F(21,170) = 1.13, p = 0.320, Adj. R² = 0.014), thus not shown in the table.

| Model | Term | Estimate | Std Error | Statistic | p value |
| --- | --- | --- | --- | --- | --- |
| Full model | | | | | |
|  | Body mass | -0.002 | 0.012 | -0.139 | 0.890 |
|  | Abs. Latitude (Range centroid) | -0.003 | 0.010 | -0.288 | 0.774 |
|  | Extent of occurrence | 0.000 | 0.011 | 0.005 | 0.996 |
|  | Web capacity index | 0.011 | 0.017 | 0.666 | 0.506 |
|  | Years known to science | 0.004 | 0.009 | 0.477 | 0.634 |
|  | Generation length | -0.009 | 0.013 | -0.704 | 0.482 |
|  | Relative beak length | -0.014 | 0.007 | -1.886 | 0.060 |
|  | Relative tail length | -0.003 | 0.007 | -0.424 | 0.672 |
|  | Use-Food [Yes] | -0.030 | 0.022 | -1.366 | 0.173 |
|  | Use-Pet [Yes] | -0.028 | 0.020 | -1.371 | 0.171 |
|  | Use-Sport [Yes] | 0.081 | 0.042 | 1.933 | 0.054 |
|  | Use-Other [Yes] | 0.007 | 0.060 | 0.122 | 0.903 |
|  | Endemic status [Yes] | -0.016 | 0.019 | -0.816 | 0.415 |
|  | Anthropogenic habitat [Yes] | 0.004 | 0.017 | 0.231 | 0.817 |
|  | Non-migratory [Yes] | 0.023 | 0.024 | 0.959 | 0.338 |
|  | Population trend [Increasing] | 0.092 | 0.038 | 2.444 | 0.015 |
|  | Population trend [Stable] | -0.040 | 0.024 | -1.676 | 0.095 |
|  | Genuine change [Yes] | 0.069 | 0.027 | 2.618 | 0.009 |
|  | BL news mentioned [Yes] | -0.070 | 0.031 | -2.299 | 0.022 |
|  | Direction [Uplist] | 0.000 | 0.023 | -0.014 | 0.989 |
| Full model (Sensitivity check removing ‘BL news mentioned’) | | | | | |
|  | Body mass | -0.002 | 0.012 | -0.193 | 0.847 |
|  | Abs. Latitude (Range centroid) | -0.003 | 0.010 | -0.272 | 0.786 |
|  | Extent of occurrence | 0.000 | 0.011 | 0.031 | 0.975 |
|  | Web capacity index | 0.021 | 0.016 | 1.301 | 0.194 |
|  | Years known to science | 0.005 | 0.009 | 0.566 | 0.572 |
|  | Generation length | -0.011 | 0.013 | -0.834 | 0.405 |
|  | Relative beak length | -0.015 | 0.007 | -2.027 | 0.043 |
|  | Relative tail length | 0.000 | 0.007 | -0.029 | 0.977 |
|  | Use-Food [Yes] | -0.025 | 0.022 | -1.157 | 0.248 |
|  | Use-Pet [Yes] | -0.025 | 0.020 | -1.233 | 0.218 |
|  | Use-Sport [Yes] | 0.056 | 0.041 | 1.388 | 0.166 |
|  | Use-Other [Yes] | -0.002 | 0.060 | -0.031 | 0.975 |
|  | Endemic status [Yes] | -0.011 | 0.019 | -0.581 | 0.562 |
|  | Anthropogenic habitat [Yes] | 0.010 | 0.017 | 0.575 | 0.566 |
|  | Non-migratory [Yes] | 0.017 | 0.024 | 0.688 | 0.492 |
|  | Population trend [Increasing] | 0.083 | 0.038 | 2.199 | 0.029 |
|  | Population trend [Stable] | -0.042 | 0.024 | -1.764 | 0.079 |
|  | Genuine change [Yes] | 0.042 | 0.024 | 1.755 | 0.080 |
|  | Direction [Uplist] | -0.006 | 0.023 | -0.255 | 0.799 |

**Table S6**. Full coefficient estimates for the genuine change interaction models. Four Linear Mixed-Effects Models testing whether the public response to Red List category changes differs between genuine and non-genuine changes. Model A is a treated-only model (N = 380 treated species, 3,420 observations) testing whether immediate and sustained effects differ by the genuineness of change among species that underwent a category change. Model B extends the primary DiD specification to include a three-way interaction with change authenticity, using the full matched dataset (N = 736 species, 6,624 observations). Models C and D repeat the treated-only analysis on the uplist-only (N = 130 treated, 82 genuine) and downlist-only (N = 192 treated, 13 genuine) matched subsets, respectively. All models include the full set of baseline covariates and random slopes (1 + time | scientific name).

| Model | Term | Estimate | Std. Error | t | p value |
| --- | --- | --- | --- | --- | --- |
| Model A: Genuine change, Treated-only LMM (N = 3420 obs, 380 species) | | | | | |
|  | Time (baseline trend) | 0.074 | 0.008 | 9.04 | <0.001*** |
|  | Body mass | 0.199 | 0.057 | 3.47 | <0.001*** |
|  | Absolute latitude | 0.219 | 0.040 | 5.55 | <0.001*** |
|  | Web Capacity Index | 0.151 | 0.048 | 3.17 | 0.002** |
|  | Years known to science | 0.197 | 0.041 | 4.78 | <0.001*** |
|  | Extent of Occurrence | 0.118 | 0.049 | 2.40 | 0.017* |
|  | Generation length | 0.188 | 0.061 | 3.05 | 0.002** |
|  | Relative beak length | 0.027 | 0.033 | 0.82 | 0.412 |
|  | Relative tail length | 0.029 | 0.033 | 0.87 | 0.387 |
|  | Use-Food | 0.208 | 0.106 | 1.97 | 0.049* |
|  | Use-Pet | 0.386 | 0.086 | 4.51 | <0.001*** |
|  | Use-Sport | 0.342 | 0.197 | 1.74 | 0.083. |
|  | Use-Other | 0.556 | 0.263 | 2.11 | 0.035* |
|  | Endemic species | 0.149 | 0.082 | 1.83 | 0.068. |
|  | Anthropogenic habitat use | -0.067 | 0.072 | -0.93 | 0.352 |
|  | Non-migrant status-Yes | -0.112 | 0.109 | -1.03 | 0.305 |
|  | Population trend-Decreasing | -0.259 | 0.094 | -2.76 | 0.006** |
|  | Population trend-Increasing | 0.149 | 0.185 | 0.81 | 0.421 |
|  | Post-event | 0.076 | 0.019 | 4.02 | <0.001*** |
|  | Genuine change | 0.118 | 0.104 | 1.13 | 0.257 |
|  | Time since event | 0.026 | 0.008 | 3.23 | 0.001** |
|  | Post-event × Genuine change | 0.042 | 0.034 | 1.24 | 0.216 |
|  | Genuine change × Time since event | -0.013 | 0.014 | -0.94 | 0.347 |
| Model B: Genuine change, Full DiD three-way interaction (N = 3420 obs, 380 species) | | | |  |  |
|  | Time (baseline trend) | 0.074 | 0.008 | 9.06 | <0.001*** |
|  | Body mass | 0.195 | 0.058 | 3.38 | <0.001*** |
|  | Absolute latitude | 0.216 | 0.040 | 5.41 | <0.001*** |
|  | Web Capacity Index | 0.153 | 0.048 | 3.20 | 0.001** |
|  | Years known to science | 0.200 | 0.041 | 4.81 | <0.001*** |
|  | Extent of Occurrence | 0.117 | 0.050 | 2.32 | 0.021* |
|  | Generation length | 0.189 | 0.062 | 3.06 | 0.002** |
|  | Relative beak length | 0.025 | 0.033 | 0.76 | 0.445 |
|  | Relative tail length | 0.031 | 0.034 | 0.93 | 0.353 |
|  | Use-Food | 0.217 | 0.106 | 2.05 | 0.041* |
|  | Use-Pet | 0.386 | 0.086 | 4.49 | <0.001*** |
|  | Use-Sport | 0.361 | 0.198 | 1.82 | 0.069. |
|  | Use-Other | 0.562 | 0.264 | 2.13 | 0.034* |
|  | Endemic species | 0.149 | 0.082 | 1.83 | 0.069. |
|  | Anthropogenic habitat use | -0.068 | 0.072 | -0.95 | 0.343 |
|  | Non-migrant status-Yes | -0.107 | 0.109 | -0.98 | 0.328 |
|  | Population trend-Decreasing | -0.234 | 0.098 | -2.39 | 0.018* |
|  | Population trend-Increasing | 0.088 | 0.194 | 0.45 | 0.650 |
|  | Post-event | 0.052 | 0.021 | 2.51 | 0.012* |
|  | Group-Treated Uplist | -0.161 | 0.148 | -1.09 | 0.276 |
|  | Genuine change | 0.328 | 0.206 | 1.59 | 0.112 |
|  | Time since event | 0.031 | 0.009 | 3.52 | <0.001*** |
|  | Post-event × Treated Uplist | 0.132 | 0.047 | 2.79 | 0.005** |
|  | Post-event × Genuine change | 0.048 | 0.062 | 0.77 | 0.440 |
|  | Treated Uplist × Genuine change | -0.134 | 0.262 | -0.51 | 0.610 |
|  | Treated Uplist × Time since event | -0.016 | 0.019 | -0.88 | 0.381 |
|  | Genuine change × Time since event | -0.038 | 0.024 | -1.56 | 0.119 |
|  | Post-event × Treated Uplist × Genuine change | -0.111 | 0.082 | -1.35 | 0.179 |
|  | Treated Uplist × Genuine change × Time since event | 0.042 | 0.032 | 1.29 | 0.196 |
| Model C: Genuine change, Uplist-only subset (N = 130 treated, 82 genuine) | | |  |  |  |
|  | Time (baseline trend) | 0.102 | 0.019 | 5.48 | <0.001*** |
|  | Body mass | 0.214 | 0.112 | 1.91 | 0.059. |
|  | Absolute latitude | 0.222 | 0.065 | 3.43 | <0.001*** |
|  | Web Capacity Index | 0.165 | 0.074 | 2.23 | 0.028* |
|  | Years known to science | 0.370 | 0.084 | 4.40 | <0.001*** |
|  | Extent of Occurrence | 0.047 | 0.094 | 0.50 | 0.616 |
|  | Generation length | 0.279 | 0.101 | 2.77 | 0.007** |
|  | Relative beak length | 0.078 | 0.059 | 1.31 | 0.194 |
|  | Relative tail length | -0.020 | 0.060 | -0.34 | 0.734 |
|  | Use-Food | 0.072 | 0.165 | 0.44 | 0.661 |
|  | Use-Pet | 0.460 | 0.134 | 3.45 | <0.001*** |
|  | Use-Sport | 0.200 | 0.236 | 0.85 | 0.399 |
|  | Use-Other | 0.493 | 0.374 | 1.32 | 0.191 |
|  | Endemic species | -0.094 | 0.172 | -0.55 | 0.585 |
|  | Anthropogenic habitat use | -0.202 | 0.128 | -1.58 | 0.117 |
|  | Non-migrant status-Yes | -0.145 | 0.176 | -0.82 | 0.412 |
|  | Population trend-Decreasing | -0.620 | 0.665 | -0.93 | 0.353 |
|  | Population trend-Increasing | -0.680 | 0.966 | -0.70 | 0.483 |
|  | Post-event | 0.143 | 0.050 | 2.87 | 0.004** |
|  | Genuine change | 0.029 | 0.180 | 0.16 | 0.870 |
|  | Time since event | -0.007 | 0.020 | -0.37 | 0.714 |
|  | Post-event × Genuine change | -0.048 | 0.061 | -0.79 | 0.429 |
|  | Genuine change × Time since event | 0.014 | 0.025 | 0.59 | 0.559 |
| Model D: Genuine change, Downlist-only subset (N = 192 treated, 13 genuine) | | |  |  |  |
|  | Time (baseline trend) | 0.065 | 0.010 | 6.77 | <0.001*** |
|  | Body mass | 0.209 | 0.075 | 2.79 | 0.006** |
|  | Absolute latitude | 0.193 | 0.053 | 3.63 | <0.001*** |
|  | Web Capacity Index | 0.074 | 0.063 | 1.18 | 0.240 |
|  | Years known to science | 0.100 | 0.057 | 1.75 | 0.081. |
|  | Extent of Occurrence | 0.014 | 0.058 | 0.24 | 0.811 |
|  | Generation length | 0.080 | 0.074 | 1.09 | 0.277 |
|  | Relative beak length | 0.104 | 0.047 | 2.20 | 0.029* |
|  | Relative tail length | 0.006 | 0.048 | 0.12 | 0.904 |
|  | Use-Food | 0.267 | 0.153 | 1.75 | 0.082. |
|  | Use-Pet | 0.403 | 0.120 | 3.36 | <0.001*** |
|  | Use-Sport | 0.251 | 0.448 | 0.56 | 0.576 |
|  | Use-Other | 0.348 | 0.384 | 0.90 | 0.367 |
|  | Endemic species | 0.125 | 0.100 | 1.25 | 0.213 |
|  | Anthropogenic habitat use | -0.060 | 0.101 | -0.59 | 0.555 |
|  | Non-migrant status-Yes | -0.134 | 0.172 | -0.78 | 0.436 |
|  | Population trend-Decreasing | -0.036 | 0.166 | -0.22 | 0.829 |
|  | Population trend-Increasing | -0.156 | 0.392 | -0.40 | 0.691 |
|  | Post-event | 0.065 | 0.023 | 2.83 | 0.005** |
|  | Genuine change | 0.589 | 0.269 | 2.19 | 0.029* |
|  | Time since event | 0.031 | 0.010 | 3.15 | 0.002** |
|  | Post-event × Genuine change | 0.072 | 0.081 | 0.89 | 0.376 |
|  | Genuine change × Time since event | -0.055 | 0.030 | -1.82 | 0.069. |

**Table S7.** Full coefficient estimates for the news mention interaction models. Three Linear Mixed-Effects Models testing whether formal communication by BirdLife International amplifies the public response to Red List category. Model A is a treated-only model (N = 380 treated species, 3,420 observations) testing whether species mentioned in BirdLife's annual Red List update showed a stronger sustained pageview response. Models B and C repeat this analysis on the uplist-only and downlist-only matched subsets, respectively. A full-DiD specification with controls is not applicable, as control species did not undergo a category change and cannot be meaningfully coded as "mentioned" for such an event. All models include the full set of baseline covariates and random slopes (1 + time | scientific name). Note that the post-event × news mention interaction term was dropped from models due to rank deficiency, as species mentioned all went through Red List category changes at the same year. Results should be interpreted with caution due to limited statistical power: only 40 treated species (10.5%) were mentioned in the annual update.

| Model | Term | Estimate | Std. Error | t | p |
| --- | --- | --- | --- | --- | --- |
| Model A: News mention, Treated-only LMM (N = 3420 obs, 380 species) | | | |  |  |
|  | Time (baseline trend) | 0.074 | 0.008 | 9.05 | <0.001*** |
|  | Body mass | 0.199 | 0.057 | 3.47 | <0.001*** |
|  | Absolute latitude | 0.234 | 0.038 | 6.11 | <0.001*** |
|  | Web Capacity Index | 0.156 | 0.047 | 3.29 | 0.001** |
|  | Years known to science | 0.207 | 0.041 | 5.07 | <0.001*** |
|  | Extent of Occurrence | 0.117 | 0.049 | 2.38 | 0.018* |
|  | Generation length | 0.183 | 0.061 | 2.98 | 0.003** |
|  | Relative beak length | 0.028 | 0.033 | 0.85 | 0.394 |
|  | Relative tail length | 0.033 | 0.033 | 1.02 | 0.310 |
|  | Use-Food | 0.227 | 0.105 | 2.16 | 0.031* |
|  | Use-Pet | 0.390 | 0.086 | 4.55 | <0.001*** |
|  | Use-Sport | 0.328 | 0.199 | 1.65 | 0.100 |
|  | Use-Other | 0.548 | 0.264 | 2.08 | 0.038* |
|  | Endemic species | 0.155 | 0.082 | 1.90 | 0.058. |
|  | Anthropogenic habitat use | -0.064 | 0.072 | -0.89 | 0.374 |
|  | Non-migrant status-Yes | -0.123 | 0.109 | -1.13 | 0.257 |
|  | Population trend-Decreasing | -0.243 | 0.094 | -2.60 | 0.010** |
|  | Population trend-Increasing | 0.163 | 0.185 | 0.88 | 0.380 |
|  | Post-event | 0.087 | 0.017 | 5.10 | <0.001*** |
|  | News mention | 0.008 | 0.045 | 0.19 | 0.850 |
|  | Time since event | 0.021 | 0.007 | 2.90 | 0.004** |
|  | News mention × Time since event | 0.011 | 0.019 | 0.58 | 0.563 |
| Model B: News mention, Uplist-only subset | |  |  |  |  |
|  | Time (baseline trend) | 0.102 | 0.019 | 5.51 | <0.001*** |
|  | Body mass | 0.213 | 0.111 | 1.91 | 0.059. |
|  | Absolute latitude | 0.221 | 0.063 | 3.50 | <0.001*** |
|  | Web Capacity Index | 0.167 | 0.073 | 2.28 | 0.025* |
|  | Years known to science | 0.372 | 0.083 | 4.47 | <0.001*** |
|  | Extent of Occurrence | 0.049 | 0.092 | 0.53 | 0.598 |
|  | Generation length | 0.275 | 0.100 | 2.77 | 0.007** |
|  | Relative beak length | 0.078 | 0.059 | 1.33 | 0.186 |
|  | Relative tail length | -0.020 | 0.058 | -0.34 | 0.735 |
|  | Use-Food | 0.080 | 0.165 | 0.49 | 0.628 |
|  | Use-Pet | 0.463 | 0.133 | 3.48 | <0.001*** |
|  | Use-Sport | 0.178 | 0.240 | 0.74 | 0.460 |
|  | Use-Other | 0.486 | 0.374 | 1.30 | 0.196 |
|  | Endemic species | -0.086 | 0.173 | -0.50 | 0.619 |
|  | Anthropogenic habitat use | -0.206 | 0.125 | -1.65 | 0.102 |
|  | Non-migrant status-Yes | -0.151 | 0.176 | -0.86 | 0.391 |
|  | Population trend-Decreasing | -0.628 | 0.653 | -0.96 | 0.338 |
|  | Population trend-Increasing | -0.698 | 0.944 | -0.74 | 0.461 |
|  | Post-event | 0.122 | 0.034 | 3.57 | <0.001*** |
|  | News mention | -0.039 | 0.060 | -0.65 | 0.516 |
|  | Time since event | -0.004 | 0.014 | -0.32 | 0.750 |
|  | News mention × Time since event | 0.025 | 0.026 | 0.96 | 0.338 |
| Model C: News mention, Downlist-only subset | |  |  |  |  |
|  | Time (baseline trend) | 0.064 | 0.010 | 6.68 | <0.001*** |
|  | Body mass | 0.208 | 0.076 | 2.75 | 0.007** |
|  | Absolute latitude | 0.218 | 0.053 | 4.13 | <0.001*** |
|  | Web Capacity Index | 0.071 | 0.063 | 1.11 | 0.268 |
|  | Years known to science | 0.121 | 0.057 | 2.13 | 0.035* |
|  | Extent of Occurrence | 0.017 | 0.058 | 0.29 | 0.769 |
|  | Generation length | 0.072 | 0.074 | 0.98 | 0.330 |
|  | Relative beak length | 0.094 | 0.048 | 1.98 | 0.050* |
|  | Relative tail length | 0.019 | 0.048 | 0.39 | 0.694 |
|  | Use-Food | 0.245 | 0.155 | 1.58 | 0.115 |
|  | Use-Pet | 0.398 | 0.121 | 3.28 | 0.001** |
|  | Use-Sport | 0.294 | 0.452 | 0.65 | 0.517 |
|  | Use-Other | 0.254 | 0.386 | 0.66 | 0.510 |
|  | Endemic species | 0.150 | 0.101 | 1.49 | 0.138 |
|  | Anthropogenic habitat use | -0.026 | 0.101 | -0.25 | 0.800 |
|  | Non-migrant status-Yes | -0.210 | 0.170 | -1.23 | 0.220 |
|  | Population trend-Decreasing | -0.026 | 0.168 | -0.15 | 0.879 |
|  | Population trend-Increasing | 0.072 | 0.410 | 0.18 | 0.861 |
|  | Post-event | 0.073 | 0.023 | 3.25 | 0.001** |
|  | News mention | -0.045 | 0.186 | -0.24 | 0.807 |
|  | Time since event | 0.026 | 0.009 | 2.75 | 0.006** |
|  | News mention × Time since event | 0.037 | 0.071 | 0.53 | 0.597 |

**Table S8.** Covariate balance before and after hierarchical matching for the Difference-in-Differences analyses

| Subset | Covariate | Treated Mean Before Matching | Control Mean Before Matching | SMD Before Matching | Treated Mean After Matching (All Pairs) | Control Mean After Matching (All Pairs) | SMD After Matching (All Pairs) |
| --- | --- | --- | --- | --- | --- | --- | --- |
| Overall |  |  |  |  |  |  |  |
|  | Body mass | 4.324 | 3.886 | 0.281 | 4.414 | 4.411 | 0.002 |
|  | Latitude (signed) | -2.040 | 2.078 | -0.220 | -1.092 | 1.327 | -0.132 |
|  | Web capacity index | 0.037 | 0.074 | -0.450 | 0.037 | 0.043 | -0.097 |
|  | Years known to science | 154.106 | 178.315 | -0.477 | 153.682 | 154.884 | -0.022 |
|  | Use-Food | 0.190 | 0.133 | 0.155 | 0.205 | 0.208 | -0.007 |
|  | Use-Pet | 0.244 | 0.395 | -0.328 | 0.268 | 0.297 | -0.064 |
|  | Use-Game | 0.033 | 0.039 | -0.030 | 0.037 | 0.042 | -0.027 |
|  | Use-Other | 0.021 | 0.020 | 0.007 | 0.018 | 0.021 | -0.019 |
|  | Population trend | 0.173 | 0.317 | -0.339 | 0.171 | 0.150 | 0.057 |
| Uplist |  |  |  |  |  |  |  |
|  | Body mass | 4.734 | 3.886 | 0.525 | 4.740 | 4.632 | 0.065 |
|  | Latitude (signed) | -1.440 | 2.078 | -0.182 | -1.702 | 0.510 | -0.119 |
|  | Web capacity index | 0.046 | 0.074 | -0.318 | 0.048 | 0.053 | -0.059 |
|  | Years known to science | 175.747 | 178.315 | -0.052 | 176.077 | 173.785 | 0.046 |
|  | Use-Food | 0.301 | 0.133 | 0.417 | 0.323 | 0.292 | 0.067 |
|  | Use-Pet | 0.377 | 0.395 | -0.038 | 0.377 | 0.400 | -0.047 |
|  | Use-Game | 0.075 | 0.039 | 0.157 | 0.085 | 0.038 | 0.193 |
|  | Use-Other | 0.041 | 0.020 | 0.124 | 0.031 | 0.038 | -0.042 |
|  | Population trend | 0.007 | 0.317 | -0.928 | 0.008 | 0.008 | 0.000 |
| Downlist |  |  |  |  |  |  |  |
|  | Body mass | 4.144 | 3.886 | 0.170 | 4.154 | 4.237 | -0.053 |
|  | Latitude (signed) | -2.303 | 2.078 | -0.237 | -0.533 | 0.032 | -0.034 |
|  | Web capacity index | 0.032 | 0.074 | -0.513 | 0.033 | 0.034 | -0.040 |
|  | Years known to science | 144.619 | 178.315 | -0.673 | 144.880 | 147.156 | -0.043 |
|  | Use-Food | 0.141 | 0.133 | 0.024 | 0.146 | 0.167 | -0.057 |
|  | Use-Pet | 0.186 | 0.395 | -0.473 | 0.229 | 0.286 | -0.131 |
|  | Use-Game | 0.015 | 0.039 | -0.148 | 0.016 | 0.031 | -0.103 |
|  | Use-Other | 0.012 | 0.020 | -0.063 | 0.016 | 0.036 | -0.131 |
|  | Population trend | 0.246 | 0.317 | -0.158 | 0.089 | 0.089 | 0.000 |

**Table S9**. Description of variables, data sources, and processing steps used in the baseline interest models

| # | Variable | Description | Type | Source | Access | Processing | N |
| --- | --- | --- | --- | --- | --- | --- | --- |
| **Response variables** | | |  |  |  |  |  |
| 1 | Wikipedia pageviews (public interest) | Total user pageviews across all Wikipedia language editions, summed over July 2015 – December 2025 | Count | Wikipedia (Wikimedia) | Wikimedia REST API (pageviews endpoint) | Wikidata ID matched via 4-round pipeline (OneZoom crosswalk → Wikidata SPARQL → English Wikipedia Search API → manual curation); per-language page titles retrieved from Wikidata API; monthly user pageviews queried and summed across languages | 10947 |
| 2 | Papers in WOK (scientific interest) | Total number of peer-reviewed articles and reviews with the species' scientific name or synonyms in the title, abstract, or keywords, published before 2026 | Count | Clarivate Web of Science | Web of Science Expanded API | Topic search (TS=) with accepted scientific name joined with all recognised synonyms via OR; server-side deduplicated count from the all-databases endpoint (databaseId=WOK) | 11185 |
| **Predictor variables** | | |  |  |  |  |  |
| 3 | Body mass | Body mass (g), log-transformed and z-scored | Continuous | AVONET (Tobias et al. 2022) | Direct download | ln-transformed, then scaled to mean = 0, SD = 1 | 10870 |
| 4 | Relative beak length | Beak length (culmen) expressed relative to body mass, accounting for phylogeny | Continuous | AVONET (Tobias et al. 2022) | Direct download | Residuals from phylogenetic regression of culmen length against ln(body mass); z-scored | 10870 |
| 5 | Relative tail length | Tail length expressed relative to body mass, accounting for phylogeny | Continuous | AVONET (Tobias et al. 2022) | Direct download | Residuals from phylogenetic regression of tail length against ln(body mass); z-scored | 10870 |
| 6 | Anthropogenic habitat | Species uses urban, suburban, or artificial terrestrial environments | Binary (Yes/No) | IUCN Red List v2025-2 | IUCN Red List API v4 | Coded Yes if any IUCN habitat classification (v3.1) record for the species includes urban, suburban, or artificial terrestrial environments | 11185 |
| 7 | Web capacity index (WCI) | Proportion of global internet users residing in countries where the species has an extant native range (reference year 2019) | Continuous | IUCN Red List; ITU ICT Indicators Database; World Bank | IUCN Red List API v4; World Bank DataBank | Country presence from IUCN range data; internet penetration rate * population per country; summed across range countries and divided by global total; missing ITU values imputed by linear interpolation then sub-region median; z-scored | 10994 |
| 8 | Years known to science | Years elapsed since formal taxonomic description (as of 2026) | Continuous | IUCN Red List v2025-2 | IUCN Red List API v4 | 2026 minus year extracted from the authority string in the IUCN taxon record; z-scored | 11185 |
| 9 | Red List category (Threatened) | Whether the species is classified as threatened under the IUCN Red List | Binary (Yes/No) | IUCN Red List v2025-2 | IUCN Red List API v4 | Most recent global assessment; VU, EN, CR -> Threatened (Yes); LC, NT -> Non-threatened (No); DD (N=36), EX, and EW (N=169) excluded | 10980 |
| 10 | Extent of occurrence (EOO) | Geographic range size (km2), log-transformed and z-scored | Continuous | IUCN Red List v2025-2 | IUCN Red List API v4 | String values parsed to numeric; ln-transformed, then z-scored | 11097 |
| 11 | Latitude (absolute range centroid) | Absolute latitudinal midpoint of the species' geographic range | Continuous | BirdLife International range maps | Provided by BirdLife International | Absolute value of the range centroid latitude; z-scored | 10974 |
| 12 | Generation length | Average age of breeding individuals (years), z-scored | Continuous | IUCN Red List v2025-2 | IUCN Red List API v4 | z-scored | 11170 |
| 13 | Non-migratory status | Species is not a migrant | Binary (Yes/No) | IUCN Red List v2025-2 | IUCN Red List API v4 | 'Not a Migrant' -> Yes; Full Migrant, Altitudinal Migrant, Nomadic -> No; species with unknown or missing movement patterns excluded (N=33) | 11174 |
| 14 | Population trend | Current direction of population change | Categorical (4 levels) | IUCN Red List v2025-2 | IUCN Red List API v4 | Four levels: Increasing, Stable (reference), Decreasing, Unknown | 11016 |
| 15 | Endemic status | Species' extant native range is confined to a single country | Binary (Yes/No) | IUCN Red List v2025-2 | IUCN Red List API v4 | IUCN sub-national and non-standard location codes resolved to ISO 3166-1 alpha-3 country codes; species present in exactly one country coded as Yes | 11185 |
| 16 | Use - Food | Species is consumed by humans | Binary (Yes/No) | IUCN Red List v2025-2 | IUCN Red List API v4 | IUCN Use & Trade classification (v1.0) code 1 | 11185 |
| 17 | Use - Pet | Species is kept or traded as a pet or for display | Binary (Yes/No) | IUCN Red List v2025-2 | IUCN Red List API v4 | IUCN Use & Trade classification (v1.0) code 13 | 11185 |
| 18 | Use - Sport | Species is hunted for sport (specimen collection also included under this IUCN code but represents a trivial fraction for birds) | Binary (Yes/No) | IUCN Red List v2025-2 | IUCN Red List API v4 | IUCN Use & Trade classification (v1.0) code 15 | 11185 |
| 19 | Use - Other | Species has other consumptive uses (e.g., medicines, handicrafts) | Binary (Yes/No) | IUCN Red List v2025-2 | IUCN Red List API v4 | Any IUCN Use & Trade classification code other than 1, 13, or 15 | 11185 |
